# A multiplexed cell-free assay to screen for antimicrobial peptides in double emulsion droplets

**DOI:** 10.1101/2021.11.17.468707

**Authors:** Nicola Nuti, Philipp Rottmann, Ariane Stucki, Philipp Koch, Sven Panke, Petra S. Dittrich

## Abstract

The global surge in bacterial resistance against traditional antibiotics triggered intensive research for novel compounds, with antimicrobial peptides (AMPs) identified as a promising candidate. Automated methods to systematically generate and screen AMPs according to their membrane preference, however, are still lacking. We introduce a novel microfluidic system for the simultaneous cell-free production and screening of AMPs for their membrane specificity. On our device, AMPs are cell-free produced within water-in-oil-in-water double emulsion droplets, generated at high frequency. Within each droplet, the peptides can interact with different classes of co-encapsulated liposomes, generating a membrane-specific fluorescent signal. The double emulsions can be incubated and observed in a hydrodynamic trapping array or analysed via flow cytometry. Our approach provides a valuable tool for the discovery and development of membrane-active antimicrobials.

## Introduction

Infectious diseases ranked consistently as the leading cause of early death throughout all human history; a trend that has only significantly changed with the discovery of antibiotics in the late 1920s[1]. These compounds ushered in an era in which most microbial infections became manageable and easy to treat. The often excessive use of antibiotics, however, resulted in the emergence of many multidrug-resistant bacterial strains which disseminate worldwide, threaten healthcare systems and cause an ever-increasing amount of deaths and economic damage[2]. For the past 20 years, drug development efforts towards novel antibiotic compounds were frequently unsuccessful, with only cyclic lipopeptides and oxazolidinones reaching the market[3]. As pointed out by the Infectious Diseases Society of America, it will be necessary to develop at least ten different antibiotic compounds within the next decade, to not face the challenges of returning to a pre-antibiotic era[4].

Receiving more and more attention since their discovery in the 1980s, antimicrobial peptides (AMPs) are promising candidates for the next generation of bactericidal compounds[5,6]. AMPs are short naturally-occurring peptides found amongst all kingdoms of life, mostly comprised of cationic and hydrophobic amino acids, capable of neutralizing a broad spectrum of pathogens[7]. Even though a small fraction of AMPs are active against intracellular targets, the vast majority of them interact with lipid membranes in a receptor-free manner and compromise their integrity, ultimately leading to cell disruption and death[8–10]. Contrary to traditional antibiotics, AMPs show the capacity to elude bacterial resistance for an extended time *in vitro*[11]. An increasing amount of *in vivo* data shows also how these molecules can coordinate an immunomodulatory response from both the innate and adaptive immune system, act as antibiofilm agents, and even neutralize several different classes of endotoxins[12–14]. To highlight the versatility of these compounds outside of their strict antimicrobial activity, they are now generally referred to as host defence peptides (HDPs), a term that better describes their pleiotropic functions within higher eukaryotes[15].

The primary molecular basis for AMP selectivity lies in their cationic nature[16]. Most AMPs show a net positive charge due to the abundance of Arg and Lys residues within their sequences and would interact preferentially with negatively charged membranes[17,18]. Contrary to mammalian cells, where negatively charged phospholipids are generally confined within the inner leaflet, bacterial membranes are rich in negatively charged phospholipids, such as phosphatidylglycerol and cardiolipin[19]. Moreover, different rates of proton exchange between mammalian and bacterial cells cause an ulterior negative shift in transmembrane potential across bacterial membranes, which will further facilitate the association of cationic peptides with the lipid bilayer[20]. After this initial electrostatic interaction, nearly all AMPs lyse the membrane via four different mechanisms of action, i.e. barrel-stave, toroidal pore, carpet, and aggregate model[16,21]. Unfortunately, many AMPs also show a certain degree of toxicity against mammalian membranes[22]. This toxicity is thought to originate from weak hydrophobic interactions between the nonpolar moieties of the peptides and the zwitterionic phospholipids commonly found on mammalian membranes, such as phosphatidylcholine[23]. Since membrane specificity is an essential feature for the development of AMPs as pharmaceutical drugs, every candidate must be screened not only for its antimicrobial activity but also for the potential activity against mammalian cells. Currently, more than 3200 AMPs have been reported in the continuously growing antimicrobial peptide database[24]. Novel peptides are found in tissue extracts of newly discovered organisms[25], identified via genome mining[26–28], or generated by engineering rationally[10,18] or systematically[29,30] known sequences by substitution with both canonical or non-canonical amino acids.

The first step to screen a library of antimicrobial peptides requires their physical production, but the traditional methods to generate peptides present several drawbacks. Purification of AMPs from their native organisms is laborious, translates in very poor yields, and it is restricted to cultivable microorganisms[31]. Chemical synthesis allows a variety of options, including the incorporation of non-canonical amino acids, but can be very expensive and time-consuming[32]. Cell-based expression of cloned AMP genes is the most economical method. The bacteriolytic nature of AMPs, however, often conflicts with the cultivation requirements of the hosts, making only a few AMPs capable of being produced this way [33].

Cell-free protein synthesis offers a promising alternative for the production of AMPs[34]. It allows for the incorporation of non-canonical amino acids and does not suffer from the toxicity of the newly produced AMPs[35]. Moreover, it does not require complicated cloning steps as it can accept linear DNA templates generated by polymerase chain reaction (PCR)[36], and it neither requires the addition of signal peptides or leader sequences, nor of dedicated transporters for the secretion of mature peptides[37]. Prokaryotic cell-free extracts, in particular, offer a very high production yield at a very low cost[38], and are frequently employed for the production of water-soluble proteins, as well as membrane proteins[39,40]. Nevertheless, they are lacking post-translational modification (PTM) capabilities, and are limited in their scope to either prokaryotic AMPs or eukaryotic AMPs that do not require them[41]. This limit can be overcome for some PTMs with site-directed incorporation of non-canonical amino acids that are chemically decorated with the required modifications[42].

Once the AMPs are synthesized and available, they need to be tested for their lipid membrane interactions. A first step, devoid of all complications arising from handling live cells in their complexity[43–47], and allowing for greater automation and scalability[48], is the *in vitro* membrane lysis assay of AMPs using artificial lipid bilayers. This is by far the most widely used method to assess the membrane disruption potential of peptide candidates, and it has been shown to correlate strongly with the *in vivo* performance of AMPs[49,50]. Membrane permeabilization is commonly measured by the efflux of a membrane-impermeant fluorescent dye from a previously loaded artificial lipid vesicle of a defined lipid composition[51].

Our primary motivation was to accelerate and automatize the process of AMP gene expression, membranolytic assay for mammalian- and bacteria-like model membranes, and sorting of the AMPs based on their membrane selectivity. With this goal in mind, we introduce a novel system and assay for the integrated cell-free production and immediate screening of AMPs for their antimicrobial activity within water-in-oil-in-water double emulsions, produced on a microfluidic device.

The use of cell-free extract in emulsion droplets for the *in vitro* screening of proteins was envisioned by Tawfik and Griffith[52], and has greatly improved directed evolution methods. Droplet compartmentalization physically links genotype and phenotype in separate pico-to nanolitre compartments, allowing for the screening of large libraries in very small volumes. These methods were further improved with the introduction of high-speed droplet microfluidic techniques coupling droplet formation with downstream analysis and sorting of individual droplets[53]. The recent development of microfluidic methods for the production of water-in-oil-in-water double emulsions provides additional advantages with respect to stability, storage and incubation, and it allows for downstream fluorescence activated cell sorting (FACS), using commercially available instruments[54]. Moreover, double emulsions have stronger stability to water evaporation and pH changes, allowing for long term incubation, and offering a stable environment for cell-free protein synthesis.

In our study, we co-encapsulate the cell-free extract, DNA templates and “sensor” liposomes in the double emulsion. Within each double emulsion droplet created with our device, a particular AMP is produced in large quantities via cell-free protein synthesis from a DNA template. The peptide can immediately interact with the membrane of co-encapsulated large unilamellar vesicles (LUVs), disrupting them and generating a fluorescent signal. As we can provide two different populations of LUVs with different lipid compositions, loaded with different spectrally separated fluorescent dyes within the same double emulsions, the dye leakage we can observe is proportional to the preference of the AMP for the specific LUV composition. By using mammalian and bacteria-like membrane compositions, we are able to screen in two separate channels for antimicrobial activity and host safety within the same double emulsion and at the same time.

## Results and Discussion

### Formation of the double emulsions

We generated monodisperse double emulsions on a microfluidic device with inlets for the inner aqueous phase, the oil phase and the outer aqueous phase, respectively (Figure 1, Supplementary Figure 1a). At first, we generate a water-in-oil emulsion that is immediately conveyed into a stream of outer aqueous phase to form a water-in-oil-in-water double emulsion (DE) droplet (Figure 1a, b).

**Figure 1.**
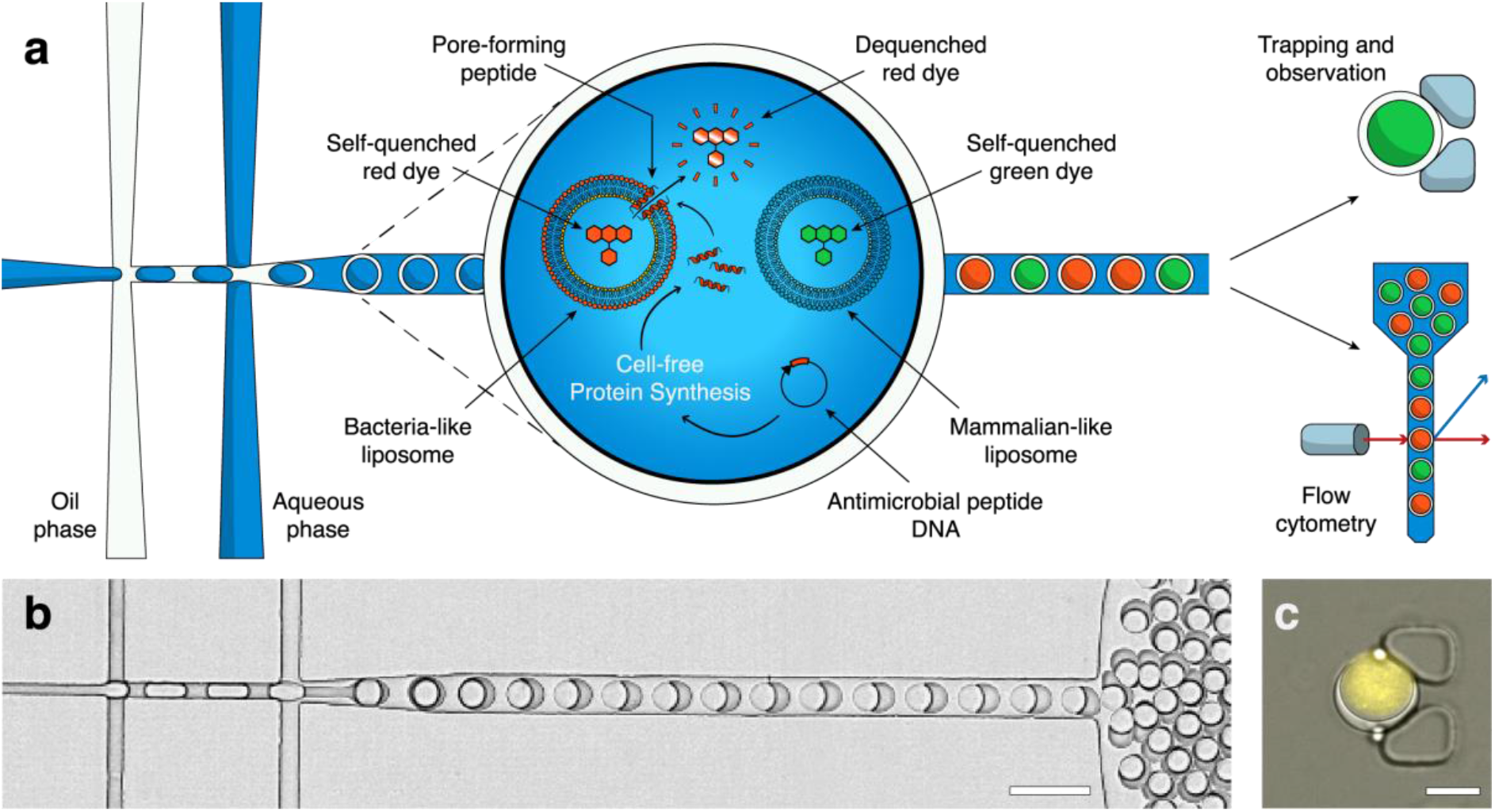
Production and analysis of antimicrobial peptides in double emulsions. (**a**) Double emulsions are formed on a microfluidic device. Within each double emulsion we express a particular AMP via cell-free protein synthesis from a DNA template. The peptide may or may not interact with the co-encapsulated LUVs, disrupting their membranes with various mechanisms. As we loaded the LUVs with a self-quenching concentration of fluorescent dye, their disruption causes release and dilution of the dye, generating a fluorescent signal. By encapsulating two different populations of LUVs with different lipid compositions (mammalian-like or bacteria-like, respectively) and loading them with different spectrally separated fluorescent dyes, we can determine antimicrobial activity and host safety simultaneously. The double emulsion droplets can then be observed over a long time in a hydrodynamic trapping array on a microfluidic device or analysed via flow cytometry. (**b**) Bright-field image of the double emulsions droplets produced on the microfluidic chip (scale bar 40 µm). (**c**) Overlaid fluorescence and bright-field image of a double emulsion in a hydrodynamic trap, containing LUVs loaded with a self-quenching concentration of SRB in the cell-free extract, showing background fluorescence (scale bar 20 µm).

Inner and outer solutions are carefully equilibrated so that the osmolarity of the outer aqueous phase matches the encapsulated inner aqueous phase. At flow rates of 1-2 µl min-1 inner aqueous phase, 1-1.5 µl min-1 oil, and 3-5 µl min-1 outer aqueous phase, we create DE droplets at a frequency of 0.3-1 kHz, with an inner aqueous droplet diameter of ∼28 µm and an outer oil shell diameter of ∼32 µm (Figure 1c).

We used the fluorinated oil HFE-7500 (3M) supplemented with 2% FluoroSurfactant (RAN Biotechnologies) to stabilize the DE droplets. After production, the DE droplets leaving the outlet are collected in an Eppendorf tube. For observation on a microscope, they can be reinjected into a second microfluidic device (Supplementary Figure 1b), where they are immobilized in a hydrodynamic trap formed by two PDMS pillars (Figure 1c). The device is conceptually similar to previously published designs for capturing single cells[55]. Alternatively, the double emulsions can be directly loaded into a flow-cytometer for high-throughput analysis, requiring no further dilution or buffer exchange.

### Stability of the assay

Fist, we confirmed the functionality of the cell-free extract, as well as its compatibility with the DE droplets, by synthesizing super-folder green fluorescent protein (sfGFP) (Figure 2a). The sfGFP plasmid (Supplementary Table 1) was mixed with the cell-free extract at an 8 nM concentration and encapsulated in the DE droplets. The DE droplets were transferred into a PDMS trapping array where they were incubated at room temperature (∼25°C) and monitored by fluorescence microscopy. The fluorescence increase indicates the protein expression until a plateau is reached after about 2 h from the addition of the plasmid.

**Table 1.**
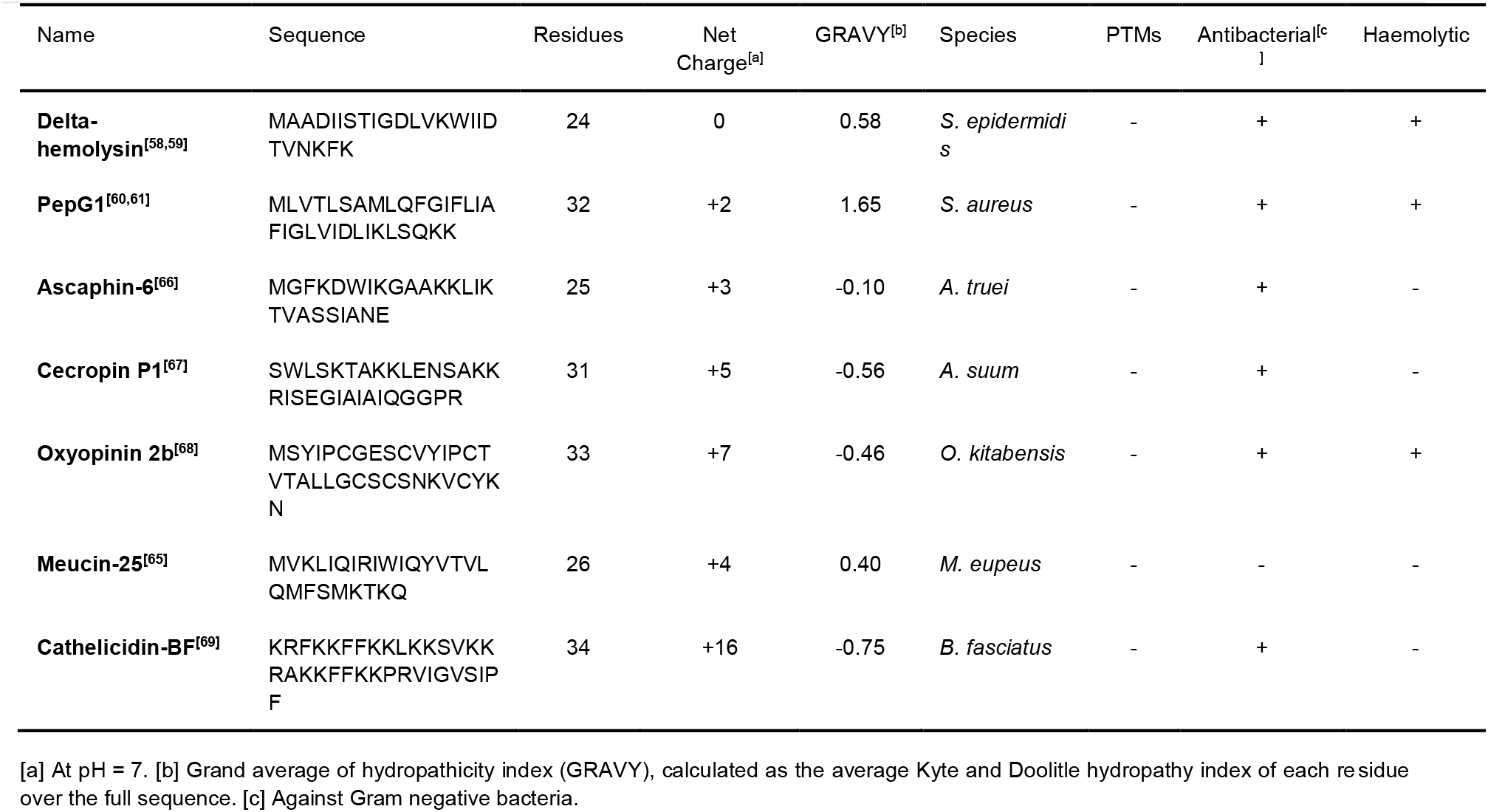
Tested antimicrobial peptides.

**Figure 2.**
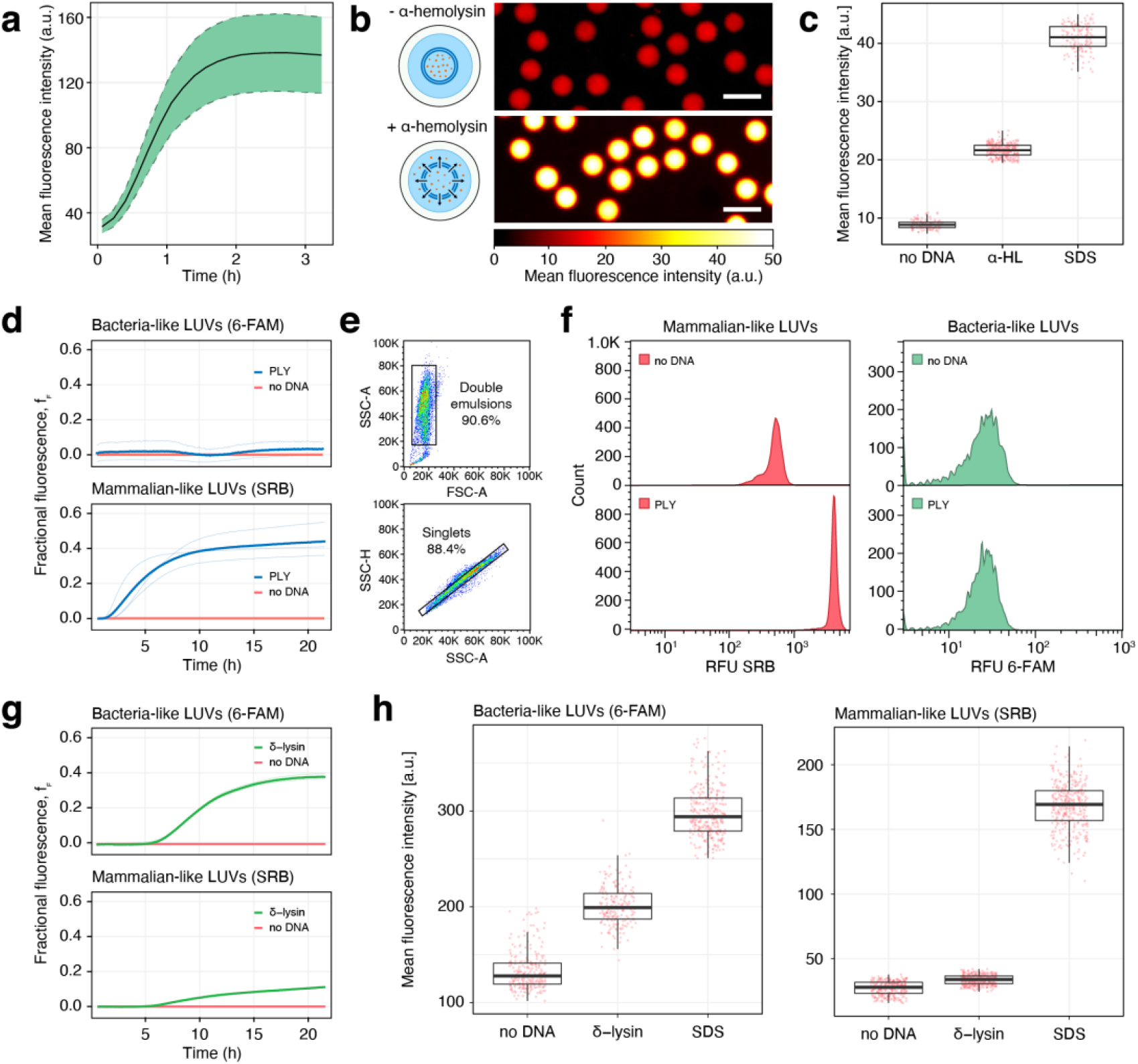
Characterization of antimicrobial peptide assay. (a) Cell-free protein production. Cell-free production of sfGFP in double emulsion (DE) droplets. The expression of sfGFP was monitored by the increase of fluorescence at 516 nm (ex. 488 nm). The dashed ribbon represents standard deviation (n = 150). **(b, c) Assay with alpha-hemolysin**. (**b**) Effect of alpha-hemolysin on mammalian-like LUVs. Fluorescence microscopy pictures of DEs containing mammalian-like LUVs loaded with a self-quenching concentration of SRB, cell-free extract, but no plasmid, showing only background fluorescence (top). With the addition of the alpha-hemolysin encoding plasmid DNA to the cell-free extract in the droplet, the DEs show a substantial increase in fluorescence due to pore formation (bottom) and the dilution of the dye beyond the self-quenching effect. Scale bars 50 µm. (**c**) Mean fluorescence intensities of b) after incubation at room temperature for 16 hours. *no DNA*: DEs without any alpha-hemolysin plasmid DNA (n = 107), *α-HL*: DEs with the alpha-hemolysin plasmid DNA (n = 258), *SDS*: double emulsions without any alpha-hemolysin plasmid DNA, exposed to a solution of 0.5% SDS in buffer throughout the incubation (n = 204). **(d-f) Assay with pneumolysin**. (**d**) Fluorophore leakage kinetic from mammalian-like LUVs with SRB and from bacteria-like LUVs with 6-FAM, induced by the cell-free expression of pneumolysin in a 384 well-plate, starting at time 0. Fractional fluorescence (*f*F) is calculated by setting the zero level to the vesicle fluorescence in the absence of DNA, and the maximum level of fluorescence, scaled to a value of 1, to the value obtained by lysing the vesicles with 0.5% SDS. Solid lines represent the average of three independent reactions visible below. (**e**) Flow cytometry plots. Top: forward scatter amplitude (FSC-A) vs. side scatter amplitude (SSC-A) and bottom: side scatter amplitude vs. side scatter height (SSC-H). Sequential gates are visible, used to select double emulsions from oil droplets, and within this population singlets from doublets. (**f**) Flow cytometry fluorescence plots of DEs containing mammalian-like LUVs with SRB and bacteria-like LUVs with 6-FAM. *no DNA*: DEs without any addition of external plasmid DNA; *PLY*: addition of the pneumolysin plasmid DNA. Fluorescence of DEs was measured after a 16-hour incubation at room temperature. **(g, h) Assay with delta-hemolysin**. (**g**) Fluorophore leakage kinetic for delta-hemolysin, conditions as in (d). (**h**) Effect of delta-hemolysin on bacteria-like LUVs and mammalian-like LUVs. Mean fluorescence intensities after incubating the DEs at room temperature for 16 hours. *no DNA*: DEs without plasmid DNA (n = 278, left, n = 261, right), *δ-lysin*: DEs with 8 nM delta-hemolysin plasmid DNA (n = 286, left, n = 310, right), *SDS*: DEs without external plasmid DNA, exposed to a solution of 0.5% SDS in buffer throughout the incubation (n = 315, left, n = 327, right). All plasmid constructs used in this Figure are shown in Supplementary Table 1.

Next, we validated the stability of LUVs in the cell-free extract within double emulsions over time. To study the effect of AMPs on mammalian and bacteria-like membranes, we tested two representative formulations of lipids (see Methods section 1.4) for their stability in the cell-free extract. In order to avoid aggregation and fusion of the LUVs, interactions between vesicles were inhibited by the addition to the lipid mixture of 4 mol % DSPE-PEG(2000). The effects of such a concentration of pegylated lipids on membrane permeabilization are negligible[56].

We loaded both LUV populations with a self-quenched concentration of either (6)-carboxyfluorescein (6-FAM) or sulforhodamine B (SRB) and added them to the cell-free extract. Special care was taken to precisely equilibrate the osmolarity of the vesicle interior with the osmolarity of the cell-free extract. Due to their dimensions, fluorophore-laden LUVs appear within the double emulsion as a homogeneous signal rather than single bright spots, facilitating detection and image analysis. When encapsulated into monodisperse double emulsions, the mixture shows a background fluorescence that is highly homogeneous between different double emulsions, indicating a uniform loading (Supplementary Figure 2a, b). This background fluorescence remains stable over 24 h, indicating complete dye retention for both combinations of LUVs formulations and fluorophores (Supplementary Figure 2c)[57]. The double emulsions’ inner diameter (inner aqueous phase) also shows size stability over 24 h (Supplementary Figure 2d).

### Characterization of the assay

To demonstrate the assay, we tested alpha-hemolysin, an α-helix pore-forming toxin from *Staphylococcus aureus*, for its activity against mammalian-like LUVs. We encapsulated LUVs containing a self-quenching concentration of SRB in double emulsions containing the cell-free extract. With the addition of 8 nM alpha-hemolysin plasmid (Supplementary Table 1) and a 16 h incubation, the pore-forming toxin is produced by the cell-free extract and forms pores in the lipid membrane, ultimately provoking dye leakage and a measurable increase in fluorescence (Figure 2b and c). Without the addition of the alpha-hemolysin plasmid, only background fluorescence is detected.

Within each double emulsion, only a limited amount of material and energy is available for cell-free production. As soon as it is consumed, cell-free protein production stops. To quantify the extent of membrane damage caused by the cell-free produced alpha-hemolysin, we constructed a positive control that allowed us to determine the highest attainable fluorescence signal using the detergent SDS. When a solution of 0.5 % SDS was added to the immobilized DE droplets, the SDS penetrated from the outside of the double emulsion through the oil shell into the inner aqueous phase, where it disrupted the LUVs.[51] The fluorescent signal reached a plateau after about 2.5 hours, indicating complete lysis (Figure 2c, Supplementary Figure 3, Supplementary Video 1). This remote lysis does not damage the double emulsion, which retains its shape and content (although with a thinned oil shell).

**Figure 3.**
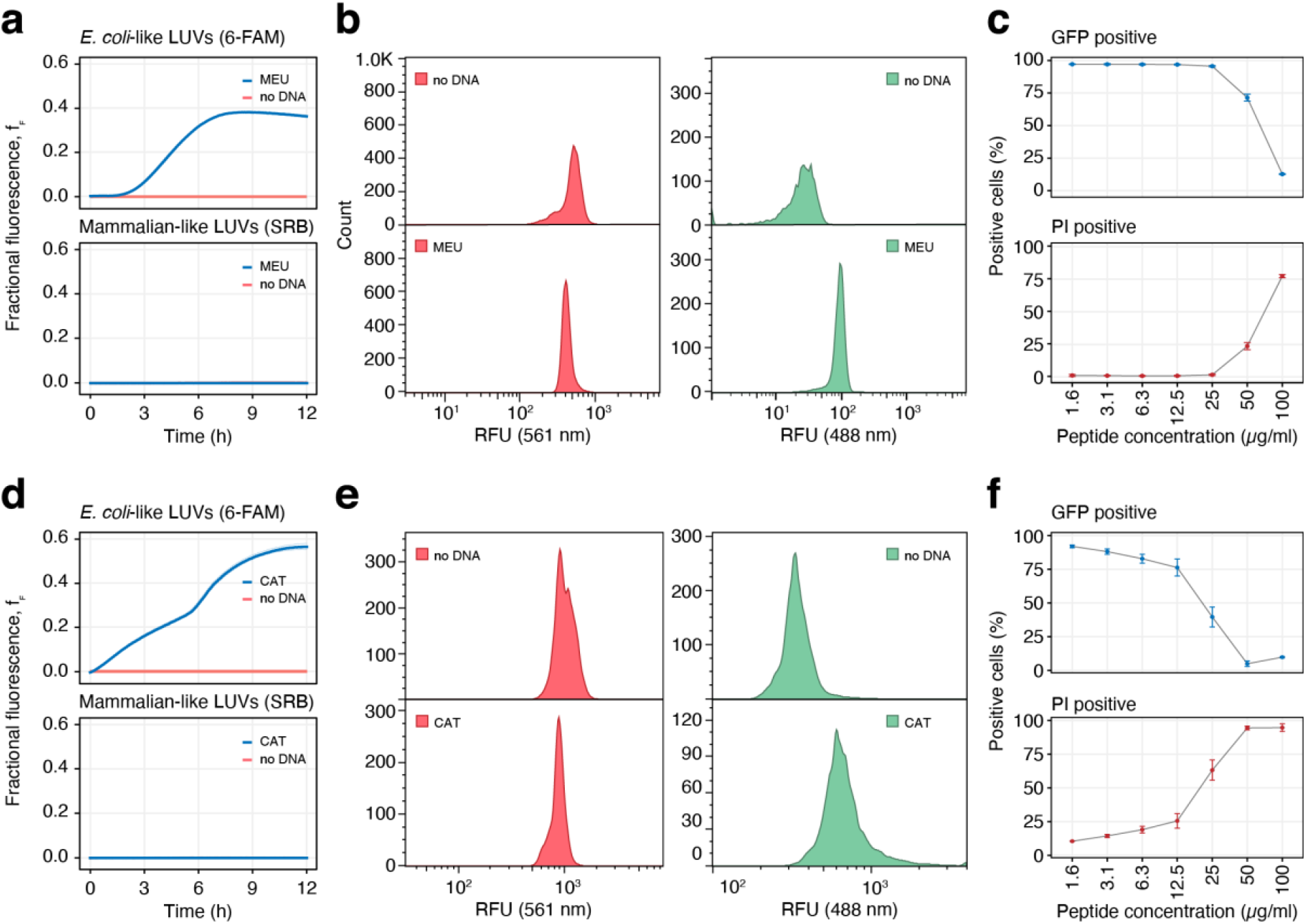
Evaluation of the antimicrobial peptides meucin-25 (a-c) and cathelicidin-BF (d-f). (**a**) Fluorophore leakage kinetic from mammalian-like LUVs with SRB and bacteria-like LUVs with 6-FAM, induced by the cell-free expression of meucin-25 in a 384 well-plate. Each well contained 8 nM of plasmid (Supplementary Table 1). Solid lines represent the average of three technical replicates displayed as well (the lines are overlapping, thus not visible). (**b**) Flow cytometry fluorescence plots of DEs containing mammalian-like LUVs with SRB, and bacteria-like LUVs with 6-FAM, measured after 16 h at RT. *no DNA*: DEs without any addition of external plasmid DNA, *MEU*: addition of 8 nM of meucin-25 plasmid. (**c**) Bacterial viability assay with increasing meucin-25 concentrations, measured by flow cytometry. Propidium iodide (PI) cannot pass intact bacterial membranes and only intercalates the DNA of permeabilized dead bacteria (“PI positive”). Constitutively expressed sfGFP protein is normally efficiently retained in intact bacterial cells (“GFP positive”) but lost in suitably permeabilized cells. Error bars indicate standard deviation (n = 10’000). (**d-f**) Fluorophore leakage kinetic (d), assay in DEs (e) and cell assays (f) for cathelicidin-BF, (Supplementary Table 1), conditions as in the corresponding figures a - c.

Pneumolysin is a β-barrel cholesterol-dependent cytolysin from *Streptococcus pneumoniae*. When the double emulsion was supplied with 8 nM of a plasmid encoding for pneumolysin (Supplementary Table 1), we could detect an increase in fluorescence due to pore formation, compared to the negative control without DNA after a 16 h incubation at room temperature (∼25°C) (Supplementary Figure 4). If we substituted the mammalian-like LUVs with bacteria-like ones, we could see no detectable increase in the same conditions. The bacteria-like LUVs, due to the absence of cholesterol, remained intact and retained their dye completely (Supplementary Figure 4).

Next, we demonstrate a multiplexed approach, where mammalian-like and bacteria-like LUVs are encapsulated together in the same DE droplets. By loading the different LUV types with two spectrally separated non-overlapping fluorescent dyes, we can detect a membrane-specific leakage as a monochrome signal. When the pneumolysin plasmid is added, the increase of fluorescence is visible for the mammalian-like LUVs only (Figure 2d, f). To show the downstream compatibility of our system with flow cytometry, we loaded the double emulsions in a flow cytometer. Based on the forward and side scatter signals we can distinguish double emulsions from oil droplets (Figure 2e), and within this population singlets (proper double emulsions) from doublets (double emulsions with two aqueous droplets within the same oil droplet). As expected, the lipid specific membrane leakage of pneumolysin can be visualized as an increase in fluorescence intensity at 561 nm, corresponding to the release of the dye encapsulated in the mammalian-like LUVs. The fluorescence intensity at 488 nm, relative to the bacteria-like LUVs, remains unchanged (Figure 2f).

As a positive control for the membrane specific disruption of the bacteria-like LUVs, we chose a delta-hemolysin variant from *Staphylococcus epidermidis* that has been shown to possess relatively broad-spectrum antimicrobial activity with limited cytotoxicity[58,59]. When the double emulsion was supplied with 8 nM of a plasmid encoding for delta-hemolysin (Supplementary Table 1), we could detect an increase in fluorescence specific for the bacteria-like membranes (Figure 2g, h). When we substituted the LUVs with mammalian-like ones, we could detect a much smaller increase in the same experimental conditions (Figure 2g, h). As an example of membrane-unspecific interaction, we finally tested PepG1, an AMP from *Staphylococcus aureus* with moderate to high haemolytic activity[60,61]. As expected, this peptide was able to permeabilize both membrane models in our assay conditions (Supplementary Figure 5).

### Screening of selected AMPs

Finally, we monitored the assay response of five additional AMPs for their antimicrobial activity and membrane specificity (Table 1).

For the peptides Ascaphin-6, Cecropin P1 and Oxyopinin 2b we did not observe a detectable interaction with the artificial membranes (Supplementary Figure 6) and hypothesized that the salt sensitivity of these peptides could be responsible for their inactivity. The cell-free extract in which LUVs and AMPs are immersed is very high in solutes, with an osmolarity more than three times higher than physiological saline solution. In such conditions, salt-sensitive peptides exhibit weaker electrostatic interactions with the lipid membrane, leading to a decrease in membranolytic activity[62-64].

As the previously tested pepG1 and delta-hemolysin had high overall hydrophobicity and were able to interact with bacteria-like membranes in our assay conditions, we expected to observe comparable activity from AMPs that presented characteristics associated with salt-resistance.

To confirm this assumption, we tested salt-resistant peptides such as meucin-25, a hydrophobic cationic peptide from the venom glands of the scorpion Mesobuthus eupeus. Meucin-25 has been shown to have potent antimalarial activity but no antibacterial or haemolytic activity at micromolar concentrations[65]. In our assay, meucin-25 did not permeabilize mammalian-like membranes but caused significant leakage from the bacteria-like liposomes (Figure 3a, b). We quantified the bacterial viability of E. coli cells exposed to different concentrations of meucin-25. The cells expressed GFP and were stained with propidium iodide (PI), a DNA-intercalating dead cell marker. If membrane damage occurs, the GFP leaks out of the cell, while PI diffuses in and intercalates the DNA. Indeed, meucin-25 showed membranolytic activity against E. coli with increasing concentrations, starting from 50 µg ml-1 (∼20 µM)(Figure 3c).

We finally tested cathelicidin-BF, an α-helical AMP found in the venom of the snake Bungarus fasciatus, which has no haemolytic activity, but a potent antimicrobial effect against Gram-negative bacteria[69]. Cathelicidin-BF has a high positive net charge and presents a linear amphipathic structure along its longitudinal axis, characteristics associated with salt-resistance[70–74]. When evaluated with our microfluidic method, cathelicidin-BF caused no dye leakage from the mammalian-like membranes and significant leakage from the bacteria-like liposomes, as expected (Figure 3d, e). The GFP-PI flow-cytometric assay for bacterial viability confirmed that there is membranolytic activity against E. coli with increasing concentrations of cathelicidin-BF, starting from as little as 12.5 µg ml-1 (∼3 µM), as shown in Figure 3f.

## Conclusion

We introduced a novel method and assay to differentiate the activity of AMPs towards specific membrane models in a microfluidic device. Our microfluidic method allows for the integrated cell-free production and screening of artificial and natural AMPs within FACS-compatible water-in-oil-in-water double emulsions, requiring only the AMP DNA sequence. We showed how AMPs interact with the membrane of co-encapsulated LUVs, generating a fluorescent readout that we can analyse via flow cytometry or on a microfluidic trapping array by time-lapse microscopy. The encapsuled 100 nm diameter sized LUVs have a high stability and can be produced in large batches, in contrast to giant unilamellar vesicles (GUVs, with a diameter over 1000 nm). Thanks to their robust dye retention, LUVs allow for the convenient detection of membrane permeabilization via the leakage and unquenching of their loaded dye. Using mammalian and bacteria-like membrane compositions, we were able to screen several AMPs for antimicrobial activity and host safety, obtaining a clear readout from salt-resistant AMPs that we could confirm using *in vivo* assays.

One of the most significant limitations of antimicrobial peptides is their salt sensitivity^[70]^. High-salt concentration environments, such as the bronchopulmonary fluids of cystic fibrosis patients, can weaken the peptide binding to the lipid bilayer, significantly reducing the efficacy of otherwise highly active antimicrobial peptides such as Human β-defensin-1, magainin 1, cecropin P1 and bactenecin^[62,63,75]^. Although the mechanism underlying salt-resistance is still poorly understood, several structural features seem to contribute to the peptide salt-resistance, such as helix stability, hydrophobicity, hydrophobic moment, net positive charge, charge distribution, and amphipathicity^[70–74]^. As the cell-free extract in which LUVs are immersed is very high in solutes, with an osmolality of ∼990 mOsm/kg (physiological solution measures ∼300 mOsm/kg), it closely mimics high-salt molecularly crowded environments, allowing for the screening of AMPs able to perform in such harsh environments.

Our novel approach simplifies the process and overcomes several limitations of previous methods for the automated screening of AMPs using artificial lipid membranes^[48]^. The assay does not require prior chemical synthesis of the peptides and relies on membranes devoid of residual solvents. Double emulsions can be prepared using simple microfluidic devices fabricated using standard soft lithography methods. Contrary to single emulsions, double emulsions are downstream compatible with traditional flow cytometry and cell sorting methods, allowing for the expression and multiplexed analysis of AMP libraries, and have stronger stability to water evaporation and pH changes, allowing for long term incubation, and offering a stable environment for cell-free protein synthesis.

Our system is open for any combination of membrane interacting polypeptides and artificial lipid vesicles, as we can easily change both LUVs’ membrane composition and co-encapsulated DNA sequences. The capacity of double emulsions to be trapped and observed long term allows for mechanistic insights on the AMP’s mode of action, offering a practical method for the screening and directed evolution of AMPs. We believe our assay will be adopted as a new tool for the pre-clinical development of the next generation of antimicrobial peptides. Finally, it is worthwhile mentioning how our system is easily adaptable for the bottom-up assembly of cell-mimicking compartments, opening up new possibilities for synthetic biology.

## Acknowledgements

Financial support from the European Research Council (Consolidator Grant HybCell, grant no. 681587) and from the Swiss National Science Foundation (NCCR Molecular Systems Engineering, project no. 51NF40-182895) is gratefully acknowledged.

