## Supplemental Information for "A multiplexed cell-free assay to screen for antimicrobial peptides in double emulsion droplets"

### Methods

#### Solutions

The buffer used for vesicle production and storage and as outer aqueous phase for all double emulsion experiments is composed of 500 mM tris(hydroxymethyl)aminomethane (TRIS HCl) pH 7.5 buffer, prepared by dissolving TRIS in Milli-Q water, followed by pH adjustment with hydrochloric acid 37%. When used as outer aqueous phase for double emulsion production, the buffer was supplemented with 2 volume % polyvinyl alcohol (PVA) (87–90% hydrolysed, molecular weight 30’000–70’000). The oil phase for double emulsion production is composed of HFE-7500 (3M) supplemented with 2 volume % FluoroSurfactant (Ran Biotech, USA). The self-quenching solutions encapsulated within the LUVs were composed of either 20 mM sulforhodamine B sodium salt in 500 mM TRIS pH 7.5, or 50 mM 5(6)-carboxyfluorescein in 500 mM TRIS pH 7.5 buffer.

#### Microfluidic chip fabrication and surface treatment

Our microfluidic setup comprises two different units, one for double emulsion production, the other for double emulsion trapping, incubation, and image acquisition. The patterned master moulds for the preparation of the chips were fabricated as follows. A 100 cm diameter silicon wafer (Si-Mat, Germany) was dehydrated for 10 min at 200 ºC, spin coated with SU-8 2015 for 30 s at 1750 rpm, and soft-baked at 95 ºC for 240 s. Using a MA6 mask aligner (Süss MicroTec, Germany), and a transparency photomask (Micro Lithography Services, UK), the wafer was exposed to a UV light dose of 150 mJ $cm^{-2}$ at 365 nm. Then, the wafer was post-baked at 95 ºC for 300 s and developed using mr-Dev 600 developer for 4 min revealing the 20 µm high features. The wafer was then rendered hydrophobic by exposing it to 1H,1H,2H,2H-perfluorodecyltrichlorosilane at 100 mbar for 12 h. The microfluidic chips were realized by pouring liquid PDMS (poly(dimethylsiloxane), oligomer to curing agent ratio of 10:1) on the silicon wafer mould to a height of 5 mm and curing it at 80 ºC for 3 h. PDMS and curing agent (Sylgard 184) were purchased from Dow Corning. After punching holes in the PDMS with a 0.5 mm inner diameter Rapid-Core biopsy puncher, the PDMS and a #1.5 glass coverslip were exposed to oxygen plasma for 45 s using a PDC-32G plasma cleaner (Harrick

Plasma), and immediately bonded to each other. The chips were then baked at 80 °C for 20 min. The production chip channel carrying the outer aqueous phase was coated with a solution of 50 mg ml$^{-1}$ polyvinyl alcohol (PVA) (87–90% hydrolysed, molecular weight 30'000–70'000, Sigma Aldrich) to increase its hydrophilicity. Positive air pressure was used to prevent the coating of the other channels, preserving their hydrophobicity. After about 3 minutes, the PVA solution was aspirated, and the chip was baked at 120°C for 20 min.

**Microfluidic chip operation for double emulsion production**

All liquids were injected in the microfluidic device using a neMESYS syringe pump (Cetoni, Germany). 1 ml Plastipak syringes (BD, Germany) were connected to the chip using PTFE microtubing (0.56 × 1.07 mm; Fisher Scientific) and metal connector tips (0.013" inner diameter, New England Small Tube). The on-chip double emulsion production was monitored using an inverted microscope equipped with a Miro M110 high-speed camera (Vision Research, New Jersey, USA). At flow rates of 1-2 µl min$^{-1}$ inner aqueous phase, 1-1.5 µl min$^{-1}$ oil, and 3-5 µl min$^{-1}$ outer aqueous phase, we created DE droplets at a frequency of 0.3-1 kHz, with an inner aqueous droplet diameter of ~28 µm and an outer oil shell diameter of ~32 µm. The inner aqueous phase consisted of cell-free extract and DNA to which the LUVs were directly added. After quantifying the LUV concentration by determining the total phospholipid content in the LUV solution, the vesicles were added to the inner aqueous phase to a final phospholipid concentration of 20 mM.

The trapping chip was mounted on an automated inverted microscope (Nikon Ti-Eclipse) and filled with buffer. The double emulsions were then flushed in the chip and got trapped in the geometric traps. While imaging, a steady flow of buffer was maintained to avoid bubble formation. Each double emulsion was imaged via brightfield and fluorescence microscopy using a 20× objective (S Plan Fluor ELWD 20×). The fluorescent excitation was performed using a Lumencor Spectra X LED light source equipped with the appropriate optical filters and dichroic mirrors (6-FAM measurement: cyan LED, 475/28 excitation filter, 495 dichroic, 525/50 emission filter; SRB measurement: green LED, 549/15 excitation filter, 562 dichroic, 593/40 emission filter). Images were recorded using a Hamamatsu Orca Flash 4 camera. The image acquisition was controlled via the NIKON NIS-Elements Advanced Research software, and the focus was kept via the Nikon Perfect Focusing System. The recorded fluorescence images were analysed using ImageJ 2.0.0 (National Institute of Health, USA).

**Vesicle preparation**

Large unilamellar vesicles were prepared with different combination of lipids and fluorescent dyes, but always following the same protocol. A total of 20 µmol of the lipids of choice were dissolved in 2 ml chloroform, and vacuum dried on the bottom of a 10 ml round glass flask using a rotary evaporator at a pressure of 200 mbar and a water bath heated to 50 °C. The lipid film was then hydrated with 0.5 ml of a solution of 20 mM SRB or 50 mM 6-FAM in TRIS pH 7.5 buffer, for 3 h under gentle swirling of the flask (orbital shaker on its lowest speed setting). The resulting solution, composed mostly of multilamellar vesicles, was frozen and thawed for 11 times in a bath of liquid nitrogen, and then in a water bath heated to 50 °C. The resulting solution, now composed of mostly unilamellar vesicles, was extruded twice using the Avanti Mini Extruder (Avanti Polar Lipids), to decrease and homogenize the size of the vesicles, and further decrease their lamellarity. The extruder was assembled with two 10 mm PE drain discs and a polycarbonate membrane with the desired pore size, both from Whatman. The lipids were extruded 11 times using first a polycarbonate membrane with

400 nm pores, and then 11 times using a membrane with 100 nm pores. The resulting solution contained both encapsulated and unencapsulated dye, and it was dialyzed using a 0.5 ml 20K MWCO Slide-A-Lyzer Dialysis Cassettes (Thermo Fisher), immersed in a 200 ml bath of 500 mM TRIS HCl pH 7.5. This was repeated three times, twice for 2 h and once overnight. The resulting solution was analysed using dynamic light scattering (DLS) on a ZetaSizer Nano (Malvern) using disposable polystyrene semimicro cuvettes (BRAND) with a path length of 1 cm (Supplementary Figure 7). 1-palmitoyl-2-oleoyl-glycero-3-phosphocholine (POPC), cholesterol, 1-palmitoyl-2-oleoyl-sn-glycero-3-phospho-(1'-rac-glycerol) (sodium salt) (POPG), 1-palmitoyl-2-oleoyl-sn-glycero-3-phosphoethanolamine (POPE), 1,2-distearoyl-sn-glycero-3-phosphoethanolamine-N-[methoxy(polyethylene glycol)-2000] (ammonium salt) (DSPE-PEG(2000)), were all purchased from Avanti Polar Lipids.

**Table 1 | LUV composition**

| | | | |
|---|---|---|---|
| **Mammalian-like** | 48% POPC | 48% Cholesterol | 4% DSPE-PEG(2000) |
| **Bacteria-like** | 72% POPE | 24% POPG | 4% DSPE-PEG(2000) |

**Cloning and plasmids**

In general, all genes for peptides were provided on high copy plasmids (pUC18 origin of replication) and expression in cell-free extract was initiated from a phage T7 promoter. Plasmids were produced in cultures with cells (such as *E. coli* Top10) which did not provide the phage T7 gene 1 (encoding T7 RNA polymerase), thus avoiding premature peptide gene expression. Plasmid construction followed standard molecular biology methods[76,77] and correct construction was confirmed by Sanger sequencing (Microsynth, Switzerland). The genes for pneumolysin and delta-hemolysin (Table 1, Supplementary Information) were chemically synthesized at IDT DNA (Belgium) and Gibson-assembled with plasmid backbone obtained by PCR (see Table 2, Supplementary Information, for primers) and plasmid pUC18_T7_sfGFP as the template. The gene for Cathelicidin-BF was synthesized by Twist-Bioscience, PCR amplified, and digested with BamHI and NdeI for ligation into the equally digested plasmid pUC18_T7_sfGFP. The genes for PepG1, Ascaphin-6, Cacropin P1, Oxyopinin 2b, and Meucin 25 were ordered with T7 promoters as plasmid constructs in pUC18 plasmid backbones from Twist Bioscience. The alpha-hemolysin encoding plasmid, which contained a theophylline riboswitch in addition to the T7 promoter, was a gift from Sheref Mansy (Addgene plasmid #53116). When used, the plasmid required the addition of 1.5 mM theophylline to activate the riboswitch. Plasmid production for cell-free protein expression was performed by overnight incubation of a 100 ml LB culture with ampicillin at 37ºC and constant agitation of 200 rpm. The plasmid was purified according to the PureLink HiPure Plasmid Midiprep Kit (Thermo Fischer Scientific) according to the manufacturer's instructions. The concentration of the purified plasmid DNA was measured with a nanodrop device (ThermoFischer Scientific).

**Cell-free production of proteins and antimicrobial peptides**

Each construct was expressed with the AccuRapid™ cell-free protein expression kit (Bioneer, South Korea). After the addition of 8 nM plasmid DNA the mixture was either encapsulated in DEs, or loaded in a 384 well plate, and incubated for at least 16 h at room temperature (~25°C). In such conditions the average concentration of protein produced after the incubation is ~0.2 mg $ml^{-1}$ (~6 µM) which was estimated based on cell-free produced sfGFP.

**Plate reader experiments**

Plate reader experiments were performed in a Cytation 5 automated plate reader (BioTek), using transparent bottom 384-well polystyrene microplates (Thermo Scientific). Within each well, the aqueous content was overlaid with 20 µl hexadecane to prevent evaporation. The plate reader was equipped with the appropriate optical filters (6-FAM/sfGFP measurement: 484/20 nm excitation filter and 530/25 nm emission filter; SRB measurement: 565/20 nm excitation filter and 606/20 nm emission filter).

Data is presented in terms of "fractional fluorescence":

$$f_F = 100 * \frac{f_{(t)} - f_{(0)}}{f_{(\infty)} - f_{(0)}}$$

Where $f_{(t)}$ is the fluorescence intensity caused by the newly cell-free produced AMP, $f_{(0)}$ is the background fluorescence intensity of the liposomes before the addition of the AMP plasmid DNA, and $f_{(\infty)}$ is the fluorescence of the LUVs completely lysed by the addition to the mixture of 0.2% Triton X-100 (for such task, Triton X-100 acts as an SDS equivalent).

**Double emulsion trapping and observation**

After assembling the microfluidic trapping device on an inverted microscope, the double emulsions were injected into the inlet port using a syringe pump at a flow rate of 1 µl $min^{-1}$. While imaging, a continuous stream of fresh buffer was kept flowing through the chamber at a rate of 0.2-0.5 µl $min^{-1}$. Each individual trap was imaged via brightfield and fluorescence microscopy, using a fully motorized inverted wide-field microscope (Nikon Ti-Eclipse). As a light source for fluorescence microscopy, we used a Lumencor Spectra X LED equipped with the appropriate optical filters and dichroic mirrors (6-FAM measurement: cyan LED, 475/28 nm excitation filter, 495 nm dichroic mirror, 525/50 nm emission filter; SRB measurement: green LED, 549/15 nm excitation filter, 562 nm dichroic mirror, 593/40 nm emission filter). Images were recorded using a Hamamatsu Orca Flash 4 camera. The microscopy setup was controlled with the Nikon NIS-Elements Advanced Research software. Fluorescence microscopy images were processed and analysed using the FIJI software and its particle analysis plugin[78]. After automatically selecting the circular regions of interest (ROIs) via thresholding, their mean grey values were measured. This process was automated using a custom FIJI script. The data was then processed using the R software. Oil droplets and overlapping double emulsions were manually omitted from the analysis.

**Flow-cytometric double emulsion analysis**

For each sample, we diluted 1 µl of double emulsions 1:200 in buffer. The sample was then analysed using an LSRFortessa™ air-cooled multi-laser benchtop flow cytometer (BD), equipped with the appropriate optical configuration (6-FAM measurements: 488 nm laser with 530/30 nm bandpass filter; SRB measurements: 561 nm laser with 586/15 bandpass filter). For each sample 10,000 events were collected. Data analysis was performed using the FlowJo software (TreeStar, USA).

**Membrane damage assay**

Chemically synthesized Meucin-25 (H-VKLIQIRIWIQYVTVLQMFSMKTKQ-OH) and Cathelicidin-BF (H- KRFKKFFKKLKKSVKKRAKKFFKKPRVIGVSIPF-OH) were purchased from Pepscan (Lelystad, NL) in >90% purity and dissolved and stored in DMSO. To measure the membrane damage of both cytoplasmic and outer membrane, we used *E. coli* TOP10 cells transformed with plasmid pSEVA 732, containing the gene for super folder green fluorescent protein (sfGFP) under the control of a strong constitutive promoter BBa_J23118 (BioBricks parts registry, parts.igem.org) and a kanamycin resistance cassette. Overnight pre-cultures were used to inoculate (1:100) fresh Mueller Hinton Broth (MHB; Sigma Aldrich) medium. Cells were grown until exponential phase (~$OD_{600}$ = 0.4), then put on ice for 20 min and finally diluted to $6 \cdot 10^5$ cells $ml^{-1}$ with MHB medium containing a final concentration of 1 µg $ml^{-1}$ propidium iodide (PI)[79]. PI is a DNA-intercalating dye, indicative of damages occurring at the cytoplasmic membrane[80]. Next, 50 µl of cell suspension were added to microtiter plate wells containing 50 µl MHB containing a 2-fold dilution series of both peptides, using 100 µg $ml^{-1}$ as the highest concentration. This was performed in duplicates. Cells and peptides were incubated at room temperature for 30 min. For each sample, the GFP and PI fluorescence of 10,000 cells was measured using a flow cytometer (LSRFortessa, BD Biosciences), equipped with the appropriate optical configuration (488 nm laser with 530/30 nm bandpass filter and 579 nm laser with 610/20 nm bandpass filter). To determine the membrane damaging properties, we calculated the percentage of GFP retaining and PI acquired cells using the FlowJo V10 software (TreeStar, USA).

**Statistical methods**

In all displayed boxplots the middle line represents the median, the boxes define the $25^{th}$ and $75^{th}$ percentiles, and the whiskers represent the standard deviation. Single data points are overlaid on the boxplot.

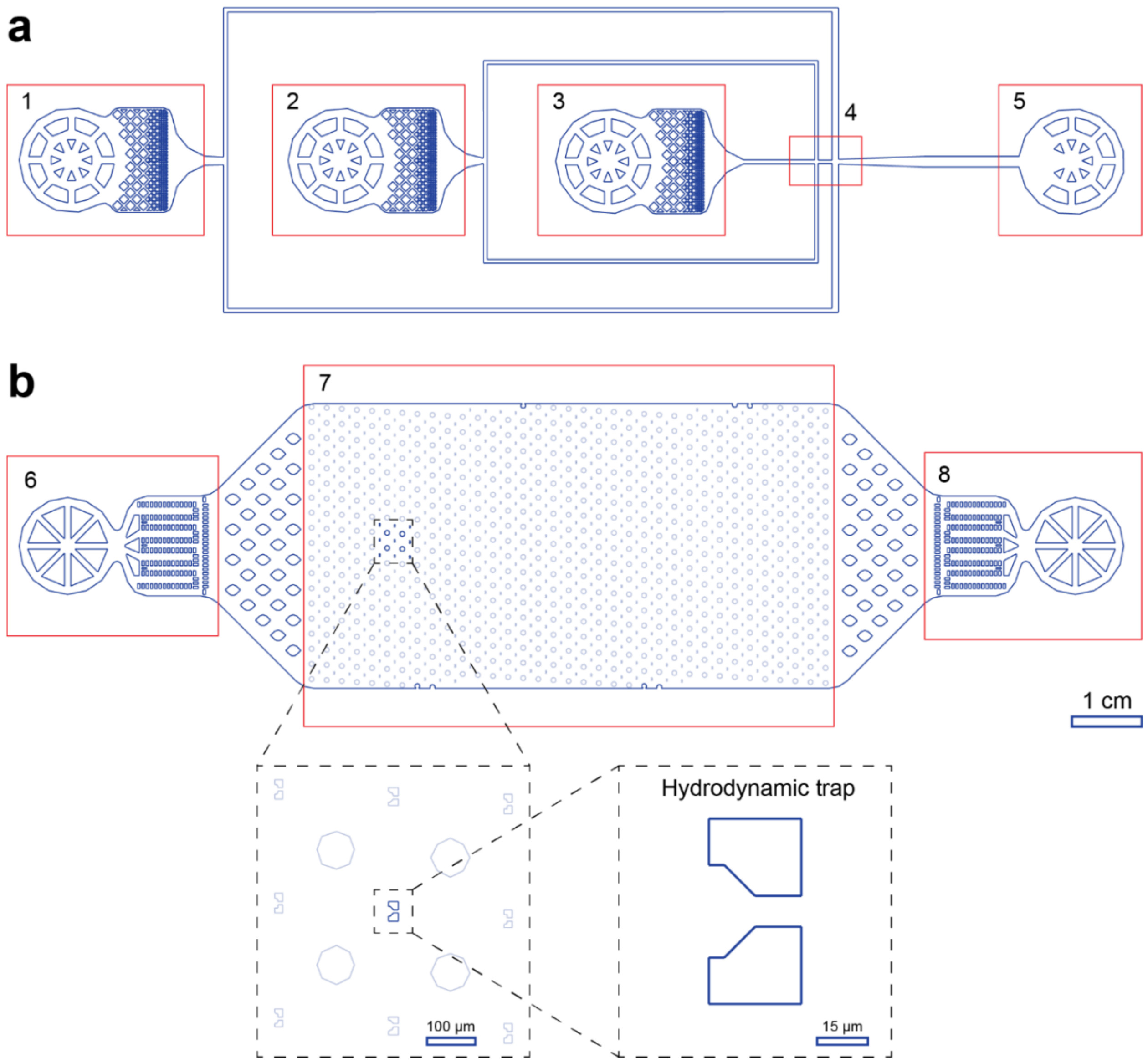


**Supplementary Figure 1 | Layout of the microfluidic devices.** (**a**) Microfluidic device for double emulsion droplet production. The device consists of three inlets for oil (2) and aqueous phases (1, 3) with filters. At the first junction (4), water-in-oil droplets are formed and immediately pushed into a stream of outer aqueous phase, forming water-in-oil-in-water double emulsion droplets. The droplets flow out of the outlet (5) and can be collected in an Eppendorf tube. (**b**) The trapping array device consists of an inlet (6) and an outlet (7), separated by a single chamber containing an array of 612 hydrodynamic traps (7). The double emulsions are physically trapped by the flow of buffer within the narrow-gapped microposts and can be released by reversing the flow direction. Both chips had structures of 20 µm in height. See also Supplementary Video 1.

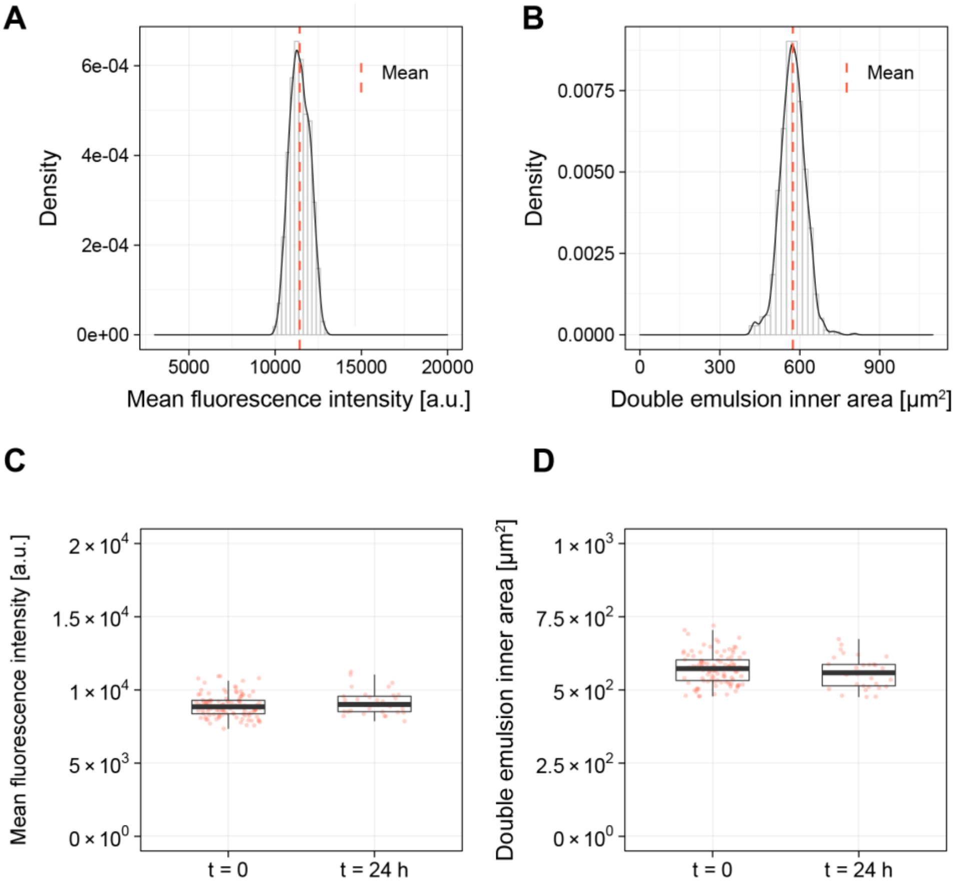


**Supplementary Figure 2 | Stability of the double emulsions containing LUVs.** (**a**) Distribution of background fluorescence between different double emulsions containing LUVs, indicating a homogeneous encapsulation (n = 1072). (**b**) Distribution of the double emulsions' inner diameter (inner aqueous phase) between different double emulsions containing LUVs, indicating a homogeneous size distribution (n = 1072). (**c**) Stability of the LUVs background fluorescence over time, indicating dye retention (n = 107 for t = 0, n = 33 for t = 24 h). (**d**) Stability of the double emulsions' inner diameter (inner aqueous phase) over time (n = 107, 33). The analysed double emulsions contained mammalian-like LUVs loaded with a self-quenching concentration of SRB.

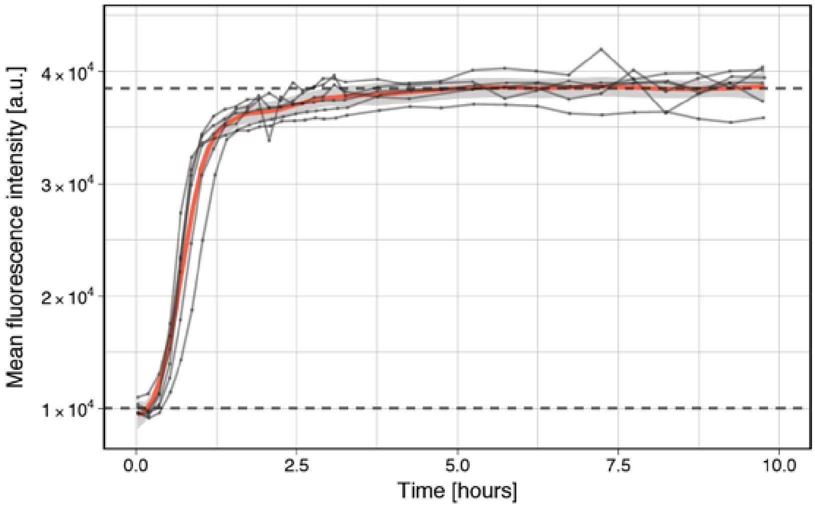


**Supplementary Figure 3 | Effect of SDS on LUVs encapsulated in double emulsions over time.** SDS penetrates through the oil shell into the inner aqueous phase of the double emulsion, where is able to lyse the fluorophore-laden LUVs. The fluorescence intensity increases and reaches a plateau when all the vesicles have been disrupted (n = 8). The vesicles are encapsulated in cell-free extract and were lysed by the addition of 0.5% SDS in buffer on the outside of the double emulsions, followed by an incubation at room temperature (~25ºC).

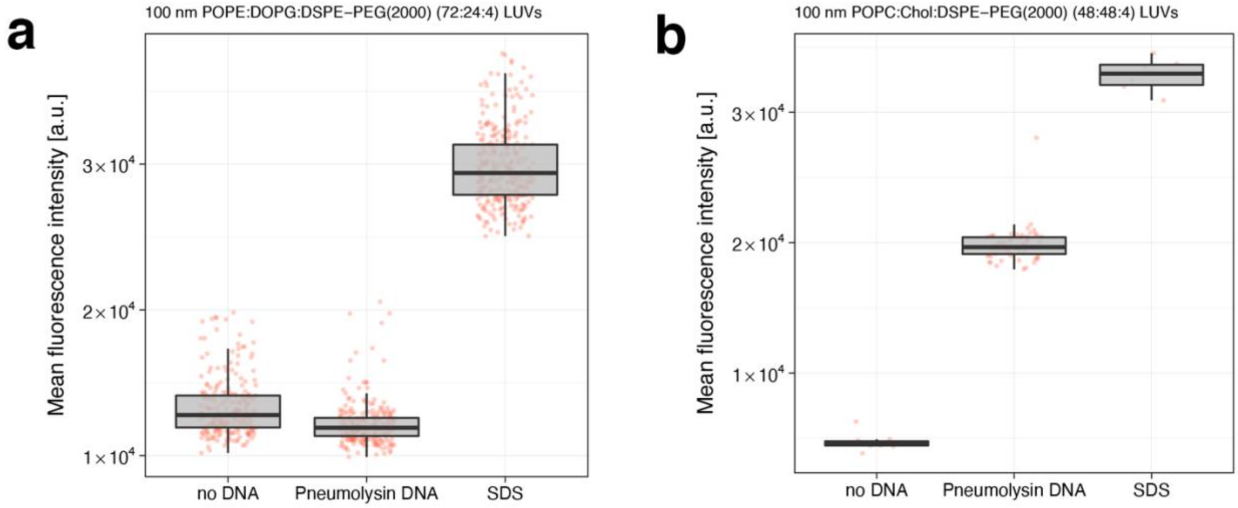


**Supplementary Figure 4 | Pneumolysin effect on different membrane compositions.** Effect of pneumolysin on bacteria-like LUVs (**a**), and mammalian-like LUVs (**b**). Mean fluorescence intensities are measured via fluorescence microscopy after incubating the double emulsions at room temperature for 16 hours. The label *no DNA* indicates double emulsions without any addition of external plasmid DNA (panel a: n = 278, panel b: n = 10), *Pneumolysin DNA* indicates double emulsions with 8 nM pneumolysin plasmid DNA (panel a: n = 267, panel b: n = 61), *SDS* indicates double emulsions without any addition of external plasmid DNA, exposed to a solution of 0.5% SDS in buffer throughout the incubation (panel a: n = 315, panel b: n = 6).

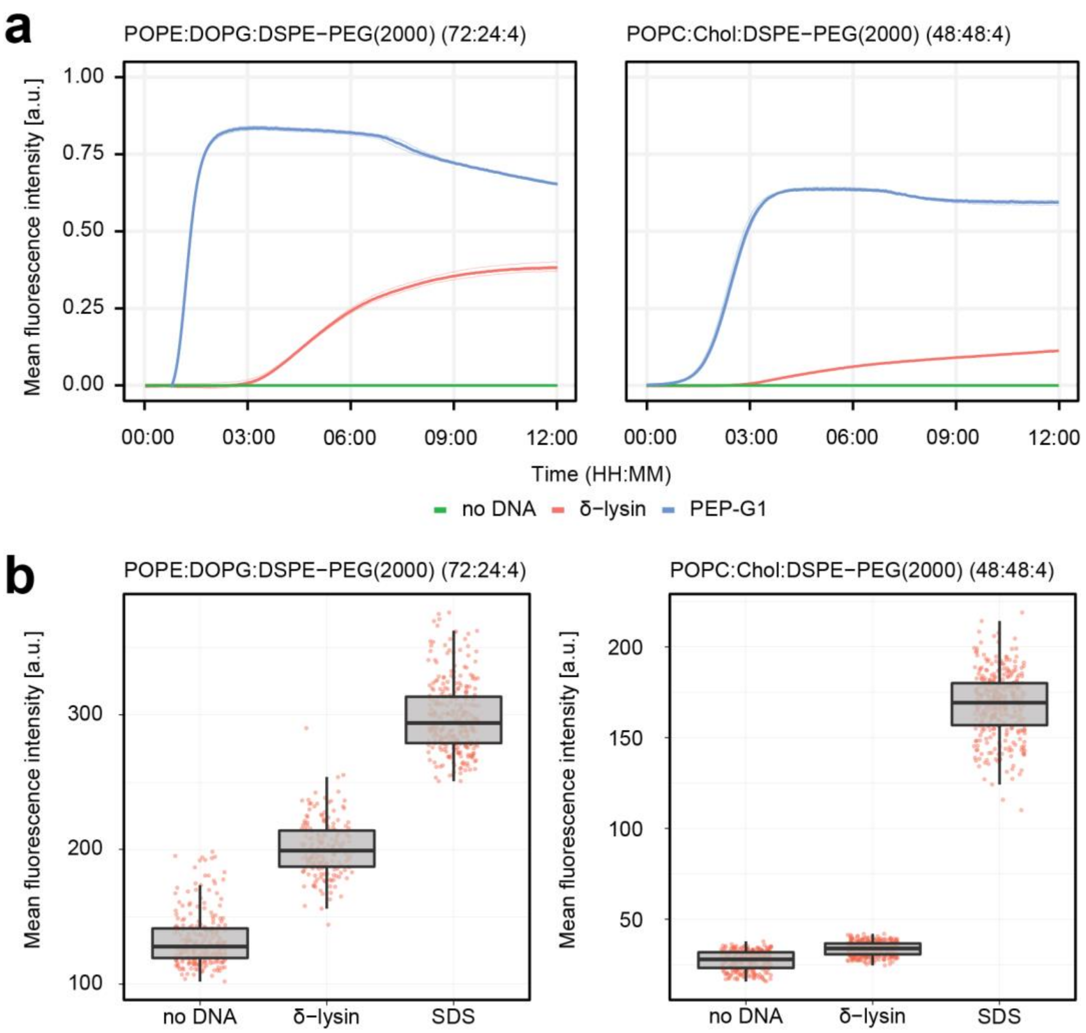


**Supplementary Figure 5 | Highly hydrophobic antimicrobial peptides interact with artificial membranes.** (**a**) Fluorophore leakage kinetic from bacteria-like LUVs loaded with a self-quenching concentration of 6-FAM (left), and mammalian-like LUVs loaded with a self-quenching concentration of SRB (right), induced by the cell-free expression of delta-hemolysin and pepG1 (sequences in Supplementary Table 1), in 384 well-plates, starting at time 0. Solid lines represent the average of three technical replicates visible below. (**b**) Effect of delta-hemolysin on bacteria-like LUVs (left), and mammalian-like LUVs (right). Mean fluorescence intensities are measured via fluorescence microscopy after incubating the double emulsions at room temperature for 16 hours. The label *no DNA* indicates double emulsions without any addition of external plasmid DNA (n = 278, left, n = 261, right), *δ-lysin* indicates double emulsions with 8 nM delta-hemolysin plasmid DNA (n = 286, left, n = 310, right), *SDS* indicates double emulsions without any addition of external plasmid DNA, exposed to a solution of 0.5% SDS in buffer throughout the incubation (n = 315, left, n = 327, right).

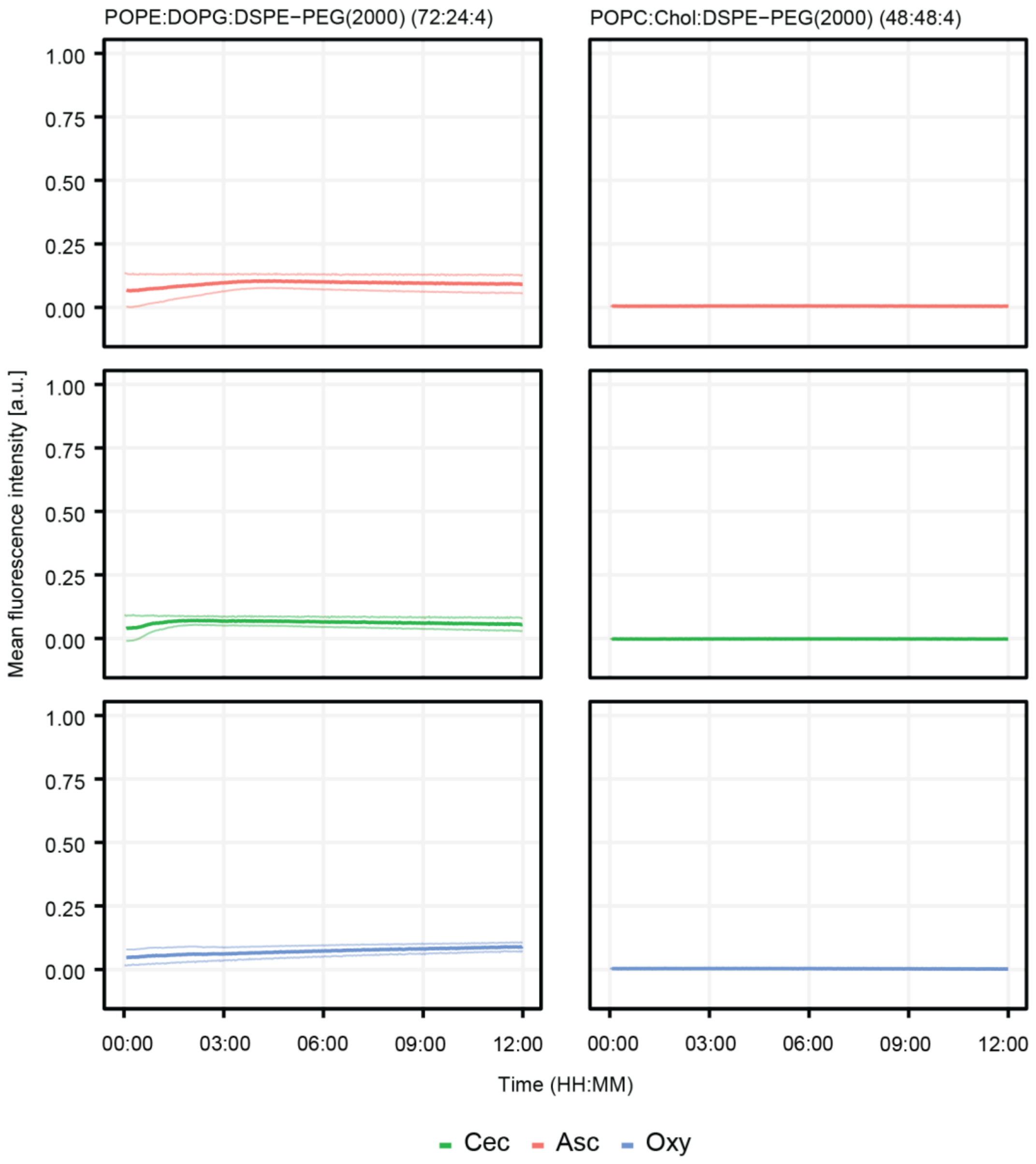


**Supplementary Figure 6 | Antimicrobial peptides with low hydrophobicity appear not to be salt-resistant.** Fluorophore leakage kinetic from bacteria-like LUVs loaded with a self-quenching concentration of 6-FAM (left), and mammalian-like LUVs loaded with a self-quenching concentration of SRB (right), induced by the cell-free expression of Cecropin P1 (Cec), Ascaphin-6 (Asc), Oxyopinin 2b (Oxy) (sequences in Supplementary Table 1), in 384 well-plates, starting at time 0. Solid lines represent the average of two independent reactions visible below. For all three experiments, alpha-hemolysin and Meucin-25 were used as positive controls.

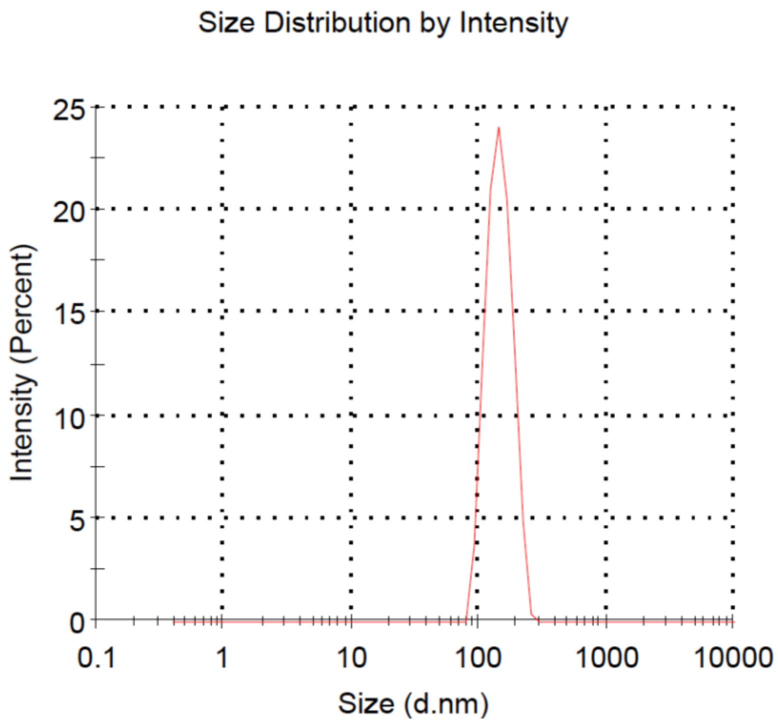


**Supplementary Figure 7 | Dynamic light scattering (DLS) size measurement of LUVs.** Representative plot for the size distribution measurement of LUVs. In the graph shown here, POPC:Chol:DSPE-PEG(2000) (48:48:4) LUVs in 350 mM Tris-HCl buffer (pH 7.5) were measured on a ZetaSizer Nano (Malvern) using disposable polystyrene semimicro cuvettes (BRAND) with a path length of 1 cm.

### 1.1 Gene fragments and primers

**Supplementary Table 1 | DNA sequences used in this work.** Start and stop codons are indicated in bold. The phage T7 promoter sequence is underlined and bold.

| Name | Sequence 5' → 3' |
|---|---|
| **Pneumolysin** | **TAATACGACTCACTATAGGG**AGACCCTCTAGAAACTGCAGTAATTTTGTTTAACTTTAAGAAGGAGATATACAT**ATG**GCAAACAAAGCAGTGAA<br>TGACTTCATTCTGGCAATGAATTATGATAAAAAAAAACTCCTGACTCATCAAGGGGAGAGTATCGAAAATCGTTTTATCAAGGAGGGTAATCAGT<br>TGCCGGACGAGTTCGTCGTCATTGAACGTAAAAAACGTTCACTGAGCACTAACACGAGCGATATTTCCGTCACGGCCACCAATGACTCACGTCTC<br>TATCCAGGGGCGCTGTTGGTGGTGGATGAAACCCTGCTGGAAAATAATCCAACCCTGCTGGCGGTGGACCGCGCACCGATGACTTATTCTATCG<br>ATTTGCCGGGCCTTGCGAGTAGCGACTCCTTTCTGCAAGTTGAAGATCCGAGCAACTCGTCCGTGCGCGGCGCCGTGAACGACCTGTTGGCCAA<br>ATGGCACCAGGACTATGGGCAGGTTAATAATGTCCCGGCGCGGATGCAATATGAGAAGATTACGGCACACTCAATGGAGCAGCTTAAAGTGAA<br>GTTCGGCTCGGACTTCGAGAAAACGGGTAACTCCTTGGACATTGATTTTAATTCAGTGCATAGTGGCGAAAAGCAAATCCAGATCGTAAATTTTA<br>AACAAATCTATTATACGGTCAGCGTTGATGCCGTGAAAAACCCGGGAGATGTGTTTCAGGATACGGTGACCGTTGAAGACCTGAAACAGCGAG<br>GTATCAGTGCGGAACGTCCACTCGTGTATATTTCCAGTGTTGCTTATGGTCGTCAAGTGTACCTCAAACTTGAAACCACCTCGAAATCTGACGAG<br>GTTGAGGCGGCATTTGAAGCCTTAATCAAAGGCGTTAAAGTGGCGCCCCAGACTGAGTGGAAACAAATCCTGGACAATACCGAAGTGAAAGCA<br>GTAATTTTAGGCGGCGATCCGAGCTCCGGCGCCCGCGTGGTTACTGGTAAGGTGGATATGGTGGAGGATCTCATTCAAGAGGGTTCACGTTTTA<br>CGGCCGATCACCCCGGGCTGCCGATTTCTTATACCACCTCTTTTTTACGCGACAATGTGGTTGCCACCTTCCAGAATAGCACTGATTACGTGGAGA<br>CCAAGGTGACTGCATATCGTAATGGAGACCTTCTTCTGGACCATAGCGGGGCCTATGTGGCGCAGTATTATATCACTTGGGATGAACTGTCGTAT<br>GATCACCAAGGTAAGGAAGTGCTGACTCCGAAGGCATGGGACCGCAATGGCCAGGACCTGACTGCGCATTTCACCACGAGTATCCCACTTAAGG<br>GTAACGTGCGTAATTTGAGCGTGAAAATTCGCGAATGCACCGGACTGGCATGGGAATGGTGGCGTACGGTTTATGAAAAGACCGATCTGCCACT<br>GGTGCGGAAACGTACCATTTCGATTTGGGGCACCACGTTATATCCCCAAGTGGAGGATAAAGTTGAAAATGATTAA**TAA**GGATCCTAGCATAAC<br>CCCTTGGGGCCTCTAAACGGGTCTTGAGGGGTTTTTTG |
| **Delta-hemolysin** | **TAATACGACTCACTATAGGG**AGACCCTCTAGAAACTGCAGTAATTTTGTTTAACTTTAAGAAGGAGATATACAT**ATG**GCGGCGGATATTATTAGC<br>ACCATTGGCGATCTGGTGAAATGGATTATTGATACCGTGAACAAATTTAAA**TAA**TTTTTCTGAAAAAATTTTAGAGTTATGTTTGATTATAGGAGA<br>ATCTGATTATTCCACGTAAGCTTTGTGCAGGGATCCTAGCATAACCCCTTGGGGCCTCTAAACGGGTCTTGAGGGGTTTTTTG |
| **Ascaphin-6** | **TAATACGACTCACTATAGGG**AGACCCTCTAGAAACTGCAGTAATTTTGTTTAACTTTAAGAAGGAGATATACAT**ATG**GGCTTTAAAGATTGGATT<br>AAAGGCGCGGCGAAAAAACTGATTAAAACCGTGGCGAGCAGCATTGCGAACGAA**TAA**TTTTTCTGAAAAAATTTTAGAGTTATGTTTGATTATA<br>GGAGAATCTGATTATTCCACGTAAGTGTGCAGGGATCCTAGCATAACCCCTTGGGGCCTCTAAACGGGTCTTGAGGGGTTTTTTG |
| **Meucin-25** | **TAATACGACTCACTATAGGG**AGACCCTCTAGAAACTGCAGTAATTTTGTTTAACTTTAAGAAGGAGATATACAT**ATG**GTGAAACTGATTCAGATT<br>CGCATTTGGATTCAGTATGTGACCGTGTTACAGATGTTTAGCATGAAAACCAAACAG**TAA**TTTTTCTGAAAAAATTTTAGAGTTATAGTGACTTAG<br>GCTATTCATAAGTTTGATTATAGGAGTGTGCAATCCTAGCATAACCCCTTGGGGCCTCTAAACGGGTCTTGAGGGGTTTTTTG |
| **Cathelicidin-BF** | **TAATACGACTCACTATAGGG**AGACCCTCTAGAAATAATTTTGTTTAACTTTAAGAAGGAGATATACAT**ATG**AAACGCTT<br>TAAAAAATTTTTTAAAAAACTGAAAAAAAGCGTGAAAAAACGCGCGAAAAAATTTTTTAAAAAACCGCGCGTGATTGG<br>CGTGAGCATTCCGTTT**TAA**TATTGATGTATCACTTACAACACGCACGTAAGCTTTGTGCAGGGATCCTAGCATAACCCC<br>TTGGGGCCTCTAAACGGGTCTTGAGGGGTTTTTTG |
| **Oxyopinin-2B** | **TAATACGACTCACTATAGGG**AGACCCTCTAGAAACTGCAGTAATTTTGTTTAACTTTAAGAAGGAGATATACAT**ATG**AGCTATATTCCGTGCGGC<br>GAAAGCTGCGTGTATATTCCGTGCACCGTGACCGCGCTGCTGGGTTGCTCGTGCAGCAACAAAGTGTGCTATAAAAAC**TAA**TTTTTCTGAAAAAA<br>TTTTAGAGTTATGTTTGATTATAGGAGTGTGCAATCCTAGCATAACCCCTTGGGGCCTCTAAACGGGTCTTGAGGGGTTTTTTG |

| | |
|---|---|
| **PepG1** | **TAATACGACTCACTATAGGG**AGACCCTCTAGAAACTGCAGTAATTTTGTTTAACTTTAAGAAGGAGATATACAT**ATG**CTGGTGACCCTGAGCGC<br>GATGTTACAGTTTGGCATTTTTCTGATTGCGTTTATTGGCCTGGTGATTGATCTGATTAAACTGAGCCAGAAAAAA**TAA**TTTTTCTGAAAAAATTTT<br>AGAGTTATGTTTGATTATAGGAGTGTGCAGGGATCCTAGCATAACCCCTTGGGGCCTCTAAACGGGTCTTGAGGGGTTTTTTG |
| **Cecropin P1** | **TAATACGACTCACTATAGGG**AGACCCTCTAGAAACTGCAGTAATTTTGTTTAACTTTAAGAAGGAGATATACAT**ATG**AGTTGGCTGAGTAAAACC<br>GCGAAAAAGTTAGAAAATAGCGCCAAAAAACGGATTAGCGAGGGTATTGCGATTGCGATTCAAGGGGGCCCTCGT**TAA**TTTTTCTGAAAAAATT<br>TTAGAGTTATAGTGACTTAGGCTATTCATAAGTTTGATTATAGGAGTGTGCAATCCTAGCATAACCCCTTGGGGCCTCTAAACGGGTCTTGAGGG<br>GTTTTTTG |
| **JF001A**<br>**(Alpha hemolysin)** | AAT**TAATACGACTCACTATAGGG**TGATACCAGCATCGTCTTGATGCCCTTGGCAGCACCCTGCTAAGGTAACAACAAG**ATG**GATTCTGATATCAA<br>TATCAAAACCGGCACCACCGATATCGGCTCCAATACCACCGTTAAAACCGGTGATCTGGTGACCTATGATAAAGAAAACGGTATGCATAAAAAA<br>GTGTTTTACTCGTTTATTGACGATAAAAACCATAACAAAAAACTGCTGGTCATCCGCACCAAAGGCACCATTGCGGGTCAATACCGTGTGTACTCC<br>GAAGAAGGTGCGAACAAAAGCGGTCTGGCTTGGCCGTCTGCCTTTAAAGTGCAGCTGCAACTGCCGGATAATGAAGTGGCGCAGATTTCAGAT<br>TATTATCCGCGTAATAGCATCGATACCAAAGAATATATGAGTACCCTGACCTATGGTTTTAATGGCAATGTTACCGGTGATGATACGGGTAAAAT<br>TGGCGGTCTGATTGGCGCCAATGTGTCCATTGGTCATACGCTGAAATACGTGCAACCGGATTTCAAAACCATTCTGGAAAGTCCGACCGATAAAA<br>AAGTGGGTTGGAAAGTTATCTTCAACAACATGGTGAATCAGAACTGGGGTCCGTACGATCGCGATTCCTGGAATCCGGTTTATGGCAATCAGCT<br>GTTTATGAAAACCCGCAACGGTAGTATGAAAGCGGCGGATAATTTTCTGGACCCGAACAAAGCCTCAAGCCTGCTGTCCAGCGGTTTTAGCCCG<br>GATTTTGCCACGGTTATTACCATGGATCGCAAAGCCAGCAAACAGCAGACCAACATTGATGTGATCTACGAACGTGTGCGTGATGATTATCAACT<br>GCATTGGACCTCAACCAATTGGAAAGGCACCAATACCAAAGATAAATGGACGGATCGCAGTTCAGAACGCTACAAAATTGATTGGGAAAAAGA<br>AGAAATGACCAAC**TAA**CTCGAGCACCACCACCACCACCACTGAGATCCGGCTGCTAACAAAGCCCGAAAGGAAGCTGAGTTGGCTGCTGCCACC<br>GCTGAGCAATAACTAGCATAACCCCTTGGGGCCTCTAAACGGGTCTTGAGGGGTTTTTTGCTGAAAGGAGGAACTATATCCGGATTGGCGAATG<br>GGACGCGCCCTGTAGCGGCGCATTAAGCGCGGCGGGTGTGGTGGTTACGCGCAGCGTGACCGCTACACTTGCCAGCGCCCTAGCGCCCGCTCC<br>TTTCGCTTTCTTCCCTTCCTTTCTCGCCACGTTCGCCGGCTTTCCCCGTCAAGCTCTAAATCGGGGGCTCCCTTTAGGGTTCCGATTTAGTGCTTTAC<br>GGCACCTCGACCCCAAAAAACTTGATTAGGGTGATGGTTCACGTAGTGGGCCATCGCCCTGATAGACGGTTTTTCGCCCTTTGACGTTGGAGTCC<br>ACGTTCTTTAATAGTGGACTCTTGTTCCAAACTGGAACAACACTCAACCCTATCTCGGTCTATTCTTTTGATTTATAAGGGATTTTGCCGATTTCGG<br>CCTATTGGTTAAAAAATGAGCTGATTTAACAAAAATTTAACGCGAATTTTAACAAAATATTAACGTTTACAATTTCAGGTGGCACTTTTCGGGGAA<br>ATGTGCGCGGAACCCCTATTTGTTTATTTTTCTAAATACATTCAAATATGTATCCGCTCATGAGACAATAACCCTGATAAATGCTTCAATAATATTG<br>AAAAAGGAAGAGTATGAGTATTCAACATTTCCGTGTCGCCCTTATTCCCTTTTTTGCGGCATTTTGCCTTCCTGTTTTTGCTCACCCAGAAACGCTG<br>GTGAAAGTAAAAGATGCTGAAGATCAGTTGGGTGCACGAGTGGGTTACATCGAACTGGATCTCAACAGCGGTAAGATCCTTGAGAGTTTTCGCC<br>CCGAAGAACGTTTTCCAATGATGAGCACTTTTAAAGTTCTGCTATGTGGCGCGGTATTATCCCGTATTGACGCCGGGCAAGAGCAACTCGGTCGC<br>CGCATACACTATTCTCAGAATGACTTGGTTGAGTACTCACCAGTCACAGAAAAGCATCTTACGGATGGCATGACAGTAAGAGAATTATGCAGTGC<br>TGCCATAACCATGAGTGATAACACTGCGGCCAACTTACTTCTGACAACGATCGGAGGACCGAAGGAGCTAACCGCTTTTTTGCACAACATGGGG<br>GATCATGTAACTCGCCTTGATCGTTGGGAACCGGAGCTGAATGAAGCCATACCAAACGACGAGCGTGACACCACGATGCCTGCAGCAATGGCAA<br>CAACGTTGCGCAAACTATTAACTGGCGAACTACTTACTCTAGCTTCCCGGCAACAATTAATAGACTGGATGGAGGCGGATAAAGTTGCAGGACC<br>ACTTCTGCGCTCGGCCCTTCCGGCTGGCTGGTTTATTGCTGATAAATCTGGAGCCGGTGAGCGTGGGTCTCGCGGTATCATTGCAGCACTGGGG<br>CCAGATGGTAAGCCCTCCCGTATCGTAGTTATCTACACGACGGGGAGTCAGGCAACTATGGATGAACGAAATAGACAGATCGCTGAGATAGGT<br>GCCTCACTGATTAAGCATTGGTAACTGTCAGACCAAGTTTACTCATATATACTTTAGATTGATTTAAAACTTCATTTTTAATTTAAAAGGATCTAGG<br>TGAAGATCCTTTTTGATAATCTCATGACCAAAATCCCTTAACGTGAGTTTTCGTTCCACTGAGCGTCAGACCCCGTAGAAAAGATCAAAGGATCTT<br>CTTGAGATCCTTTTTTTCTGCGCGTAATCTGCTGCTTGCAAACAAAAAAACCACCGCTACCAGCGGTGGTTTGTTTGCCGGATCAAGAGCTACCAA<br>CTCTTTTTCCGAAGGTAACTGGCTTCAGCAGAGCGCAGATACCAAATACTGTCCTTCTAGTGTAGCCGTAGTTAGGCCACCACTTCAAGAACTCTG<br>TAGCACCGCCTACATACCTCGCTCTGCTAATCCTGTTACCAGTGGCTGCTGCCAGTGGCGATAAGTCGTGTCTTACCGGGTTGGACTCAAGACGA<br>TAGTTACCGGATAAGGCGCAGCGGTCGGGCTGAACGGGGGGTTCGTGCACACAGCCCAGCTTGGAGCGAACGACCTACACCGAACTGAGATAC<br>CTACAGCGTGAGCTATGAGAAAGCGCCACGCTTCCCGAAGGGAGAAAGGCGGACAGGTATCCGGTAAGCGGCAGGGTCGGAACAGGAGAGC<br>GCACGAGGGAGCTTCCAGGGGGAAACGCCTGGTATCTTTATAGTCCTGTCGGGTTTCGCCACCTCTGACTTGAGCGTCGATTTTTGTGATGCTCG<br>TCAGGGGGGCGGAGCCTATGGAAAAACGCCAGCAACGCGGCCTTTTTACGGTTCCTGGCCTTTTGCTGGCCTTTTGCTCACATGTTCTTTCCTGC<br>GTTATCCCCTGATTCTGTGGATAACCGTATTACCGCCTTTGAGTGAGCTGATACCGCTCGCCGCAGCCGAACGACCGAGCGCAGCGAGTCAGTG<br>AGCGAGGAAGCGGAAGAGCGCCTGATGCGGTATTTTCTCCTTACGCATCTGTGCGGTATTTCACACCGCATATATGGTGCACTCTCAGTACAATC<br>TGCTCTGATGCCGCATAGTTAAGCCAGTATACACTCCGCTATCGCTACGTGACTGGGTCATGGCTGCGCCCCGACACCCGCCAACACCCGCTGAC<br>GCGCCCTGACGGGCTTGTCTGCTCCCGGCATCCGCTTACAGACAAGCTGTGACCGTCTCCGGGAGCTGCATGTGTCAGAGGTTTTCACCGTCATC<br>ACCGAAACGCGCGAGGCAGCTGCGGTAAAGCTCATCAGCGTGGTCGTGAAGCGATTCACAGATGTCTGCCTGTTCATCCGCGTCCAGCTCGTTG<br>AGTTTCTCCAGAAGCGTTAATGTCTGGCTTCTGATAAAGCGGGCCATGTTAAGGGCGGTTTTTTCCTGTTTGGTCACTGATGCCTCCGTGTAAGG<br>GGGATTTCTGTTCATGGGGGTAATGATACCGATGAAACGAGAGAGGATGCTCACGATACGGGTTACTGATGATGAACATGCCCGGTTACTGGA<br>ACGTTGTGAGGGTAAACAACTGGCGGTATGGATGCGGCGGGACCAGAGAAAAATCACTCAGGGTCAATGCCAGCGCTTCGTTAATACAGATGT<br>AGGTGTTCCACAGGGTAGCCAGCAGCATCCTGCGATGCAGATCCGGAACATAATGGTGCAGGGCGCTGACTTCCGCGTTTCCAGACTTTACGAA<br>ACACGGAAACCGAAGACCATTCATGTTGTTGCTCAGGTCGCAGACGTTTTGCAGCAGCAGTCGCTTCACGTTCGCTCGCGTATCGGTGATTCATT<br>CTGCTAACCAGTAAGGCAACCCCGCCAGCCTAGCCGGGTCCTCAACGACAGGAGCACGATCATGCGCACCCGTGGGGCCGCCATGCCGGCGAT<br>AATGGCCTGCTTCTCGCCGAAACGTTTGGTGGCGGGACCAGTGACGAAGGCTTGAGCGAGGGCGTGCAAGATTCCGAATACCGCAAGCGACAG<br>GCCGATCATCGTCGCGCTCCAGCGAAAGCGGTCCTCGCCGAAAATGACCCAGAGCGCTGCCGGCACCTGTCCTACGAGTTGCATGATAAAGAAG<br>ACAGTCATAAGTGCGGCGACGATAGTCATGCCCCGCGCCCACCGGAAGGAGCTGACTGGGTTGAAGGCTCTCAAGGGCATCGGTCGAGATCCC<br>GGTGCCTAATGAGTGAGCTAACTTACATTAATTGCGTTGCGCTCACTGCCCGCTTTCCAGTCGGGAAACCTGTCGTGCCAGCTGCATTAATGAAT<br>CGGCCAACGCGCGGGGAGAGGCGGTTTGCGTATTGGGCGCCAGGGTGGTTTTTCTTTTCACCAGTGAGACGGGCAACAGCTGATTGCCCTTCAC<br>CGCCTGGCCCTGAGAGAGTTGCAGCAAGCGGTCCACGCTGGTTTGCCCCAGCAGGCGAAAATCCTGTTTGATGGTGGTTAACGGCGGGATATA<br>ACATGAGCTGTCTTCGGTATCGTCGTATCCCACTACCGAGATATCCGCACCAACGCGCAGCCCGGACTCGGTAATGGCGCGCATTGCGCCCAGC<br>GCCATCTGATCGTTGGCAACCAGCATCGCAGTGGGAACGATGCCCTCATTCAGCATTTGCATGGTTTGTTGAAAACCGGACATGGCACTCCAGTC<br>GCCTTCCCGTTCCGCTATCGGCTGAATTTGATTGCGAGTGAGATATTTATGCCAGCCAGCCAGACGCAGACGCGCCGAGACAGAACTTAATGGG<br>CCCGCTAACAGCGCGATTTGCTGGTGACCCAATGCGACCAGATGCTCCACGCCCAGTCGCGTACCGTCTTCATGGGAGAAAATAATACTGTTGAT<br>GGGTGTCTGGTCAGAGACATCAAGAAATAACGCCGGAACATTAGTGCAGGCAGCTTCCACAGCAATGGCATCCTGGTCATCCAGCGGATAGTTA<br>ATGATCAGCCCACTGACGCGTTGCGCGAGAAGATTGTGCACCGCCGCTTTACAGGCTTCGACGCCGCTTCGTTCTACCATCGACACCACCACGCT<br>GGCACCCAGTTGATCGGCGCGAGATTTAATCGCCGCGACAATTTGCGACGGCGCGTGCAGGGCCAGACTGGAGGTGGCAACGCCAATCAGCAA<br>CGACTGTTTGCCCGCCAGTTGTTGTGCCACGCGGTTGGGAATGTAATTCAGCTCCGCCATCGCCGCTTCCACTTTTTCCCGCGTTTTCGCAGAAAC<br>GTGGCTGGCCTGGTTCACCACGCGGGAAACGGTCTGATAAGAGACACCGGCATACTCTGCGACATCGTATAACGTTACTGGTTTCACATTCACCA<br>CCCTGAATTGACTCTCTTCCGGGCGCTATCATGCCATACCGCGAAAGGTTTTGCGCCATTCGATGGTGTCCGGGATCTCGACGCTCTCCCTTATGC<br>GACTCCTGCATTAGGAAGCAGCCCAGTAGTAGGTTGAGGCCGTTGAGCACCGCCGCCGCAAGGAATGGTGCATGCAAGGAGATGGCGCCCAAC<br>AGTCCCCCGGCCACGGGGCCTGCCACCATACCCACGCCGAAACAAGCGCTCATGAGCCCGAAGTGGCGAGCCCGATCTTCCCCATCGGTGATGT<br>CGGCGATATAGGCGCCAGCAACCGCACCTGTGGCGCCGGTGATGCCGGCCACGATGCGTCCGGCGTAGAGGATCGAGATCTCGATCCCGCGA |
| **pUC18_T7_sfGFP** | AACGACGGCCAGTGCCAAGCTTGCGAAAT**TAATACGACTCACTATAGGG**AGACCCTCTAGAAACTGCAGTAATTTTGTTTAACTTTAAGAAGGA<br>GATATACAT**ATG**AGCAAAGGAGAAGAACTTTTCACTGGAGTTGTCCCAATTCTTGTTGAATTAGATGGTGATGTTAATGGGCACAAATTTTCTGT<br>CCGTGGAGAGGGTGAAGGTGATGCTACAAACGGAAAACTCACCCTTAAATTTATTTGCACTACTGGAAAACTACCTGTTCCGTGGCCAACACTTG<br>TCACTACTCTGACCTATGGTGTTCAATGCTTTTCCCGTTATCCGGATCACATGAAACGGCATGACTTTTTCAAGAGTGCCATGCCCGAAGGTTATG<br>TACAGGAACGCACTATATCTTTCAAAGATGACGGGACCTACAAGACGCGTGCTGAAGTCAAGTTTGAAGGTGATACCCTTGTTAATCGTATCGAG<br>TTAAAGGGTATTGATTTTAAAGAAGATGGAAACATTCTTGGACACAAACTCGAGTACAACTTTAACTCACACAATGTATACATCACGGCAGACAA<br>ACAAAAGAATGGAATCAAAGCTAACTTCAAAATTCGCCACAACGTTGAAGATGGTTCCGTTCAACTAGCAGACCATTATCAACAAAATACTCCAA<br>TTGGCGATGGCCCTGTCCTTTTACCAGACAACCATTACCTGTCGACACAATCTGTCCTTTCGAAAGATCCCAACGAAAAGCGTGACCACATGGTCC |

| | |
|---|---|
| | TTCTTGAGTTTGTAACTGCTGCTGGGATTACACATGGCATGGATGAGCTCTACAAATAATAAGGATCCTAGCATAACCCCTTGGGGCCTCTAAAC<br>GGGTCTTGAGGGGTTTTTTGAATTCGTAATCATGGTCATAGCTGTTTCCTGTGTGAAATTGTTATCCGCTCACAATTCCACACAACATACGAGCCG<br>GAAGCATAAAGTGTAAAGCCTGGGGTGCCTAATGAGTGAGCTAACTCACATTAATTGCGTTGCGCTCACTGCCCGCTTTCCAGTCGGGAAACCT<br>GTCGTGCCAGCTGCATTAATGAATCGGCCAACGCGCGGGGAGAGGCGGTTTGCGTATTGGGCGCTCTTCCGCTTCCTCGCTCACTGACTCGCTGC<br>GCTCGGTCGTTCGGCTGCGGCGAGCGGTATCAGCTCACTCAAAGGCGGTAATACGGTTATCCACAGAATCAGGGGATAACGCAGGAAAGAACA<br>TGTGAGCAAAAGGCCAGCAAAAGGCCAGGAACCGTAAAAAGGCCGCGTTGCTGGCGTTTTTCCATAGGCTCCGCCCCCCTGACGAGCATCACAA<br>AAATCGACGCTCAAGTCAGAGGTGGCGAAACCCGACAGGACTATAAAGATACCAGGCGTTTCCCCCTGGAAGCTCCCTCGTGCGCTCTCCTGTTC<br>CGACCCTGCCGCTTACCGGATACCTGTCCGCCTTTCTCCCTTCGGGAAGCGTGGCGCTTTCTCAAAGCTCACGCTGTAGGTATCTCAGTTCGGTGT<br>AGGTCGTTCGCTCCAAGCTGGGCTGTGTGCACGAACCCCCCGTTCAGCCCGACCGCTGCGCCTTATCCGGTAACTATCGTCTTGAGTCCAACCCG<br>GTAAGACACGACTTATCGCCACTGGCAGCAGCCACTGGTAACAGGATTAGCAGAGCGAGGTATGTAGGCGGTGCTACAGAGTTCTTGAAGTGG<br>TGGCCTAACTACGGCTACACTAGAAGAACAGTATTTGGTATCTGCGCTCTGCTGAAGCCAGTTACCTTCGGAAAAAGAGTTGGTAGCTCTTGATC<br>CGGCAAACAAACCACCGCTGGTAGCGGTGGTTTTTTTGTTTGCAAGCAGCAGATTACGCGCAGAAAAAAAGGATCTCAAGAAGATCCTTTGATC<br>TTTTCTACGGGGTCTGACGCTCAGTGGAACGAAAACTCACGTTAAGGGATTTTGGTCATGAGATTATCAAAAAGGATCTTCACCTAGATCCTTTTA<br>AATTAAAAATGAAGTTTTAAATCAATCTAAAGTATATATGAGTAAACTTGGTCTGACAGTTACCAATGCTTAATCAGTGAGGCACCTATCTCAGCG<br>ATCTGTCTATTTCGTTCATCCATAGTTGCCTGACTCCCCGTCGTGTAGATAACTACGATACGGGAGGGCTTACCATCTGGCCCCAGTGCTGCAATG<br>ATACCGCGAGACCCACGCTCACCGGCTCCAGATTTATCAGCAATAAACCAGCCAGCCGGAAGGGCCGAGCGCAGAAGTGGTCCTGCAACTTTAT<br>CCGCCTCCATCCAGTCTATTAATTGTTGCCGGGAAGCTAGAGTAAGTAGTTCGCCAGTTAATAGTTTGCGCAACGTTGTTGCCATTGCTACAGGC<br>ATCGTGGTGTCACGCTCGTCGTTTGGTATGGCTTCATTCAGCTCCGGTTCCCAACGATCAAGGCGAGTTACATGATCCCCCATGTTGTGCAAAAA<br>AGCGGTTAGCTCCTTCGGTCCTCCGATCGTTGTCAGAAGTAAGTTGGCCGCAGTGTTATCACTCATGGTTATGGCAGCACTGCATAATTCTCTTAC<br>TGTCATGCCATCCGTAAGATGCTTTTCTGTGACTGGTGAGTACTCAACCAAGTCATTCTGAGAATAGTGTATGCGGCGACCGAGTTGCTCTTGCCC<br>GGCGTCAATACGGGATAATACCGCGCCACATAGCAGAACTTTAAAAGTGCTCATCATTGGAAAACGTTCTTCGGGGCGAAAACTCTCAAGGATC<br>TTACCGCTGTTGAGATCCAGTTCGATGTAACCCACTCGTGCACCCAACTGATCTTCAGCATCTTTTACTTTCACCAGCGTTTCTGGGTGAGCAAAA<br>ACAGGAAGGCAAAATGCCGCAAAAAAGGGAATAAGGGCGACACGGAAATGTTGAATACTCATACTCTTCCTTTTTCAATATTATTGAAGCATTTA<br>TCAGGGTTATTGTCTCATGAGCGGATACATATTTGAATGTATTTAGAAAAA**TAA**ACAAATAGGGGTTCCGCGCACATTTCCCCGAAAAGTGCCAC<br>CTGACGTCTAAGAAACCATTATTATCATGACATTAACCTATAAAAATAGGCGTATCACGAGGCCCTTTCGTCTCGCGCGTTTCGGTGATGACGGT<br>GAAAACCTCTGACACATGCAGCTCCCGGAGACGGTCACAGCTTGTCTGTAAGCGGATGCCGGGAGCAGACAAGCCCGTCAGGGCGCGTCAGCG<br>GGTGTTGGCGGGTGTCGGGGCTGGCTTAACTATGCGGCATCAGAGCAGATTGTACTGAGAGTGCACCATATGCGGTGTGAAATACCGCACAGA<br>TGCGTAAGGAGAAAATACCGCATCAGGCGCCATTCGCCATTCAGGCTGCGCAACTGTTGGGAAGGGCGATCGGTGCGGGCCTCTTCGCTATTAC<br>GCCAGCTGGCGAAAGGGGGATGTGCTGCAAGGCGATTAAGTTGGGTAACGCCAGGGTTTTCCCAGTCACGACGTTGTAA |

**Supplementary Table 2 |** Primer list for plasmids construction of delta-hemolysin and pneumolysin.

| Name | Sequence 5’ → 3’ | Description |
|---|---|---|
| **oPR146** | GATATACATATGGCGGCGGATATTATTAG | Fwd. primer amplification delta-hemolysin gene fragment |
| **oPR147** | GGATCCCCTAGGGCGCACGTGGAATAATCAGATTCTC | Rev. primer amplification of delta-hemolysin gene fragment |
| **oPR148** | GGTCATATGGCAAACAAAGC | Fwd. primer amplification of pneumolysin gene fragment |
| **oPR149** | TGCGGATCCTTATTAATCATTTT | Rev. primer amplification of pneumolysin gene fragment |
| **oPR15** | CTTTGTTTGCCATATGTATATCTCC | Rev. primer for amplification of pUC18_T7 backbone fragment based on the pUC18_T7_sfGFP plasmid and Gibson assembly with the pneumolysin gene fragment |
| **oPR152** | TAATATCCGCCGCCATATGTATATCT | Rev. primer for amplification of pUC18_T7 backbone fragment based on the pUC18_T7_sfGFP plasmid and Gibson assembly with the delta-hemolysin gene fragment |
| **oPR160** | TGATTAATAAGGATCCTAGCATAAC | Fwd. primer for amplification of pUC18_T7 backbone fragment based on the pUC18_T7_sfGFP plasmid and Gibson assembly with the pneumolysin gene fragment |
| **oPR161** | CTAGGGGATCCTAGCATAAC | Fwd. primer for amplification of pUC18_T7 backbone fragment based on the pUC18_T7_sfGFP plasmid and Gibson assembly with the delta-hemolysin gene fragment |
| **oPR222** | ATGGGATCCCTGCACAAAGCTTACGTG | Rev. primer amplification of cathelicidin-BF gene fragment |
| **oPR221** | GCCCATATGAAACGCTTTAAAAAATTTTTTAAAAAACTG | Fwd. primer amplification of cathelicidin-BF gene fragment |
